# ABC-transporter CFTR folds with high fidelity through a modular, stepwise pathway

**DOI:** 10.1101/2022.07.20.500765

**Authors:** Jisu Im, Tamara Hillenaar, Hui Ying Yeoh, Priyanka Sahasrabudhe, Marjolein Mijnders, Marcel van Willigen, Peter van der Sluijs, Ineke Braakman

**Affiliations:** Cellular Protein Chemistry, Bijvoet Centre for Biomolecular Research, Faculty of Science, Science for Life, Utrecht University, 3584 CH Utrecht, The Netherlands

**Keywords:** Protein folding, domain assembly, COPII, secretory pathway, ABC-transporter, cystic fibrosis

## Abstract

The question how proteins fold is especially pointed for large multidomain, multispanning membrane proteins with complex topologies. We have uncovered the sequence of events that encompass proper folding of the ABC transporter CFTR in live cells, by combining kinetic radiolabeling with protease-susceptibility assays. We found that CFTR folds in two clearly distinct stages. The first, co-translational, stage involves folding of the 2 transmembrane domains TMD1 and TMD2, plus one nucleotide-binding domain, NBD1. The second stage is a simultaneous, post-translational increase in protease resistance for both TMDs and NBD2 caused by assembly of these domains onto NBD1.

Our technology probes every 2-3 residues (on average) in CFTR. This in-depth analysis at amino-acid level allows detailed analysis of domain folding and importantly also the next level: the assembly of the domains to native, folded CFTR. Defects and changes brought about by medicines, chaperones or mutations also are amenable to analysis. We here show that the DXD motif in NBD1 that was identified to be required for export of CFTR from the ER turned out to be required for proper domain folding and assembly instead, upstream of transport. CFTR mutated in this motif phenocopies the misfolding and degradation of the well-known disease-causing mutant F508del that established cystic fibrosis as protein-folding disease. The highly modular process of domain folding and stepwise domain assembly explains the relatively high fidelity of folding and the importance of a step-wise folding process for such complex proteins.

## INTRODUCTION

Membrane proteins account for one third of all of the proteins encoded by the genome of sequenced species. They mediate and integrate fundamental processes occurring on both sides of biological membranes. In humans at least 40% of membrane proteins span the membrane more than once [1]. These so called polytopic membrane proteins include amongst others receptors, transporters, and channels, and are the target of more than half of all small-molecule drugs [1, 2]. Many inherited diseases are associated with impaired folding and function of membrane proteins, signifying the importance of uncovering mechanisms underlying their folding for human health and drug development [3].

Folding of polytopic membrane proteins in vivo can be viewed as a series of sequential, overlapping steps [4]. While the nascent chain is translated by ER-associated ribosomes, transmembrane helices (TMHs) are inserted and integrated into the ER membrane. Helical packing within the membrane occurs, along with folding of soluble domains and their organization into a functional protein. Co-translational folding intermediates have been characterized for ribosome-bound nascent chains [5, 6] and domains in a few multi-domain membrane proteins have been found to fold individually and co-translationally [5, 7, 8]. Yet, how transmembrane domains fold and assemble into a mature, functional structure is still largely unknown.

We have addressed this question using the ABC transporter CFTR (cystic fibrosis transmembrane conductance regulator) as a model protein. The ABC transporter superfamiliy is one of the oldest and most conserved protein superfamilies [9, 10]. ABC transporters are multi-domain, multi-spanning membrane proteins, which transport various substrates across membranes, regulated by ATP hydrolysis, and are therefore crucial for homeostasis. CFTR functions as a chloride channel in the plasma membrane [11], and mutations in CFTR cause the disease cystic fibrosis (CF) [12].

CFTR contains two similar halves with each a transmembrane domain (TMD) and a nucleotide-binding domain (NBD). The halves are connected via the intrinsically unstructured Regulatory (R) region. The cryo-EM structures and various structural models of CFTR show that the TMDs are assembled via their transmembrane helices and helical intracellular loops (ICLs) [13-17]. The end of the ICLs, the so-called coupling helices, connect the TMDs with the NBDs. Each TMD contains two ICLs, one of which interacts with the NBD in the same half of the molecule whereas the other ICL binds to the NBD in the other half, in a cross-over fashion.

Seminal studies uncovered the orientation and integration of CFTR transmembrane (TM) helices into the membrane [18]. The signal recognition particle recognizes signal(s) in CFTR TMs and targets them to the translocon, for insertion into the ER membrane. The polypeptide sequence around transmembrane helices of polytopic proteins can cause translocation stop or re-start to allow formation of cytosolic and lumenal domains during translation. This would suggest that the first TMH initiates translocation and the next one causes stop-transfer, which continues in alternating fashion with the following TMDs. Although intuitively appealing, it does not seem to hold as common principle [19, 20]. In CFTR the first TMH only interacts with Sec61 for ∼25% of the time, while the second TMH is efficiently recognized [21]. After failed recognition, the first TMH then must be inserted into the ER-membrane following recognition of TMH2. Which one of the translocon(s) is responsible for CFTR insertion and translocation into the ER membrane however remains to be defined. The nucleotide-binding domains and the R region fold in the cytosol. The individual domains of CFTR fold mostly co-translationally, and the TMDs have been found to continue folding post-translationally, which require interactions with the other domains [7, 21]. Taken together, this supports a view that individual domains of CFTR fold mostly during translation whereas acquisition of compactly folded domains continues post-translationally through domain-domain interactions [22].

Despite this progress, mechanistic insight in the folding pathways of CFTR domains and especially of the later assembly events into a functional structure is only understood in rather general terms. This is in part because TM helices are embedded in a membrane and shielded from the aqueous environment but is also caused by the paucity of suitable reagents to study them. We here have developed and characterized antibodies against the TMDs of CFTR. We then used them together with antibodies against the NBDs to establish an integrated workflow of radiolabeling pulse chase, limited proteolysis, and immunoprecipitation, for temporal analysis of CFTR (domain) folding in vivo. By identifying the boundaries of domain-specific proteolytic fragments, we uncovered detailed changes in conformation during maturation of newly synthesized CFTR and describe a global folding profile of the CFTR protein.

## RESULTS

### Characterization of antibodies against CFTR TMDs

Limited proteolysis in combination with immune-based detection of fragments is a versatile method to assay protein folding in vivo [7, 23, 24]. Most antibodies that have been available for these studies recognize exposed cytoplasmic epitopes in CFTR, which largely precludes analysis of the transmembrane domains. To extend folding studies to the transmembrane domains of CFTR, we raised antibodies in rabbits against the small first extracellular loop, peptide aa S364-K381 of TMD1, and to residues Q1035-S1049 and E1172-Q1186 of TMD2, and named them E1-22, TMD1C, I4N, and TMD2C, respectively (Figure 1a).

**Figure 1.**
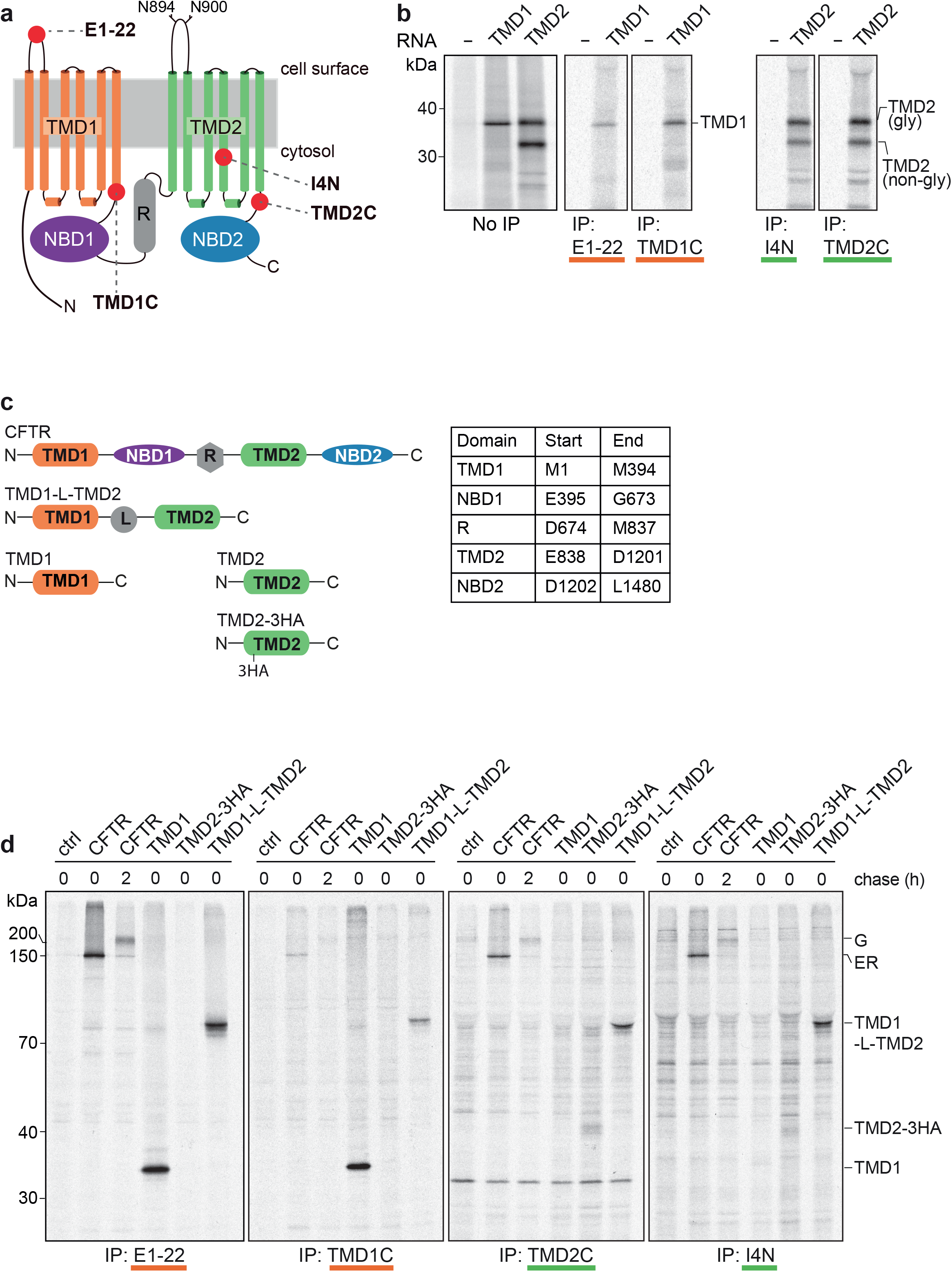
New antibodies against CFTR transmembrane domains recognize whole domains synthesized in vitro and in vivo. **(a)** Cartoon of CFTR in which red spheres represent the position of the epitopes to which antibodies were raised. Sites for N-linked glycosylation in TMD2 are shown. **(b)** TMD1 and TMD2 were translated in vitro and translocated in the presence of semi-intact cells as a source of ER membranes. Membrane fractions of in-vitro translated TMD1 and TMD2 were resolved by SDS-PAGE (left panel) or immunoprecipitated (IP) with E1-22 & TMD1C (middle), and I4N & TMD2C, respectively (right). Samples were resolved by 12% SDS-PAGE. **(c)** Schematic representation of constructs used in **(b)** and **(d)**, with a table showing the used boundaries of CFTR domains. **(d)** HEK293T cells expressing single or multi-domain CFTR constructs were pulse labeled for 15 minutes and lysed immediately or after a chase of 2 hours. CFTR was immunoprecipitated from detergent lysates with indicated antibodies and resolved by 10% SDS-PAGE.

We first tested the antibodies on TMD1 and TMD2 translated in vitro in the presence of ^35^S-labeled amino acids and a source of ER membranes (Figure 1b). TMD1 is detectable as one major protein of 35 kDa while TMD2 is resolved in bands at 32 kDa and 37 kDa. The larger form represents core-glycosylated TMD2, and the 32-kDa band is non-glycosylated TMD2 (Figure S1a; [25]. E1-22 and TMD1C immunoprecipitated TMD1 and not TMD2, while I4N and TMD2C detected both forms of TMD2 but not TMD1 (Figure 1b and not shown), showing that the antibodies recognized their target proteins in a relatively non-complex biological system, which lacks additional radiolabeled proteins.

We next asked whether the antibodies against the TMDs recognized their epitopes in intact cells. Cells expressing various single and multi-domain constructs of CFTR (Figure 1c) were labeled with ^35^S-methionine/cysteine and subsequently chased without radiolabel to follow maturation of the labeled proteins with time. Newly synthesized full-length CFTR is core glycosylated and appears at the end of a 15-min pulse as a ∼150 kDa band (Figure S1b, MrPink, band ER) [25]. During the chase with unlabeled methionine and cysteine (2 hours), CFTR is transported to the Golgi complex where the glycans are modified to complex glycans, with concomitant increase of molecular weight to ∼200 kDa (Figure S1, MrPink, band Golgi) [25].

All four antibodies recognized full-length CFTR, both the ER and Golgi forms (Figure 1d), albeit with much lower yield for TMD1C and the TMD2 antibodies (I4N, TMD2-C). Only E1-22 detected full-length CFTR as well as control antibody MrPink (cf. Figure 1d with Figures S1b and S2). Cells expressing single-domain and multi-domain constructs of CFTR were not subjected to a chase period, because these are retained in the ER and therefore do not reach the oligosaccharide-modifying enzymes of the Golgi complex [26]. E1-22 and TMD1C immunoprecipitated TMD1 linked to TMD2 and TMD1 expressed on its own (Figure 1d), similar to in-vitro translated TMD1 (Figure 1b). TMD2C and I4N detected individually expressed TMD2 but only when it contained a triple HA tag in ECL4 or was linked to TMD1 (Figure 1d), suggesting that TMD2 expressed on its own was not stable enough to allow accumulation of a detectable quantity. All four antibodies clearly retained their specificity in a radiolabeled cell lysate. TMD1C showed increased recognition of isolated TMD1 compared to TMD1 in CFTR, suggesting shielding of the TMD1C epitope in full-length CFTR.

### Antibodies reveal folding intermediates of CFTR

We next deployed the TMD antibodies in conjunction with antibodies against the two NBDs to zoom in on the folding of individual CFTR domains. We used MrPink for NBD1 and monoclonal antibody 596 for NBD2 [23, 27]. The flexible R region was omitted from the analysis because of its phosphorylation-dependent conformation [28]. After a radiolabel pulse-chase, CFTR was immunoprecipitated directly upon lysis or after subjecting the lysates to limited proteolysis using Proteinase K (Figure 2a) [7, 29].

**Figure 2.**
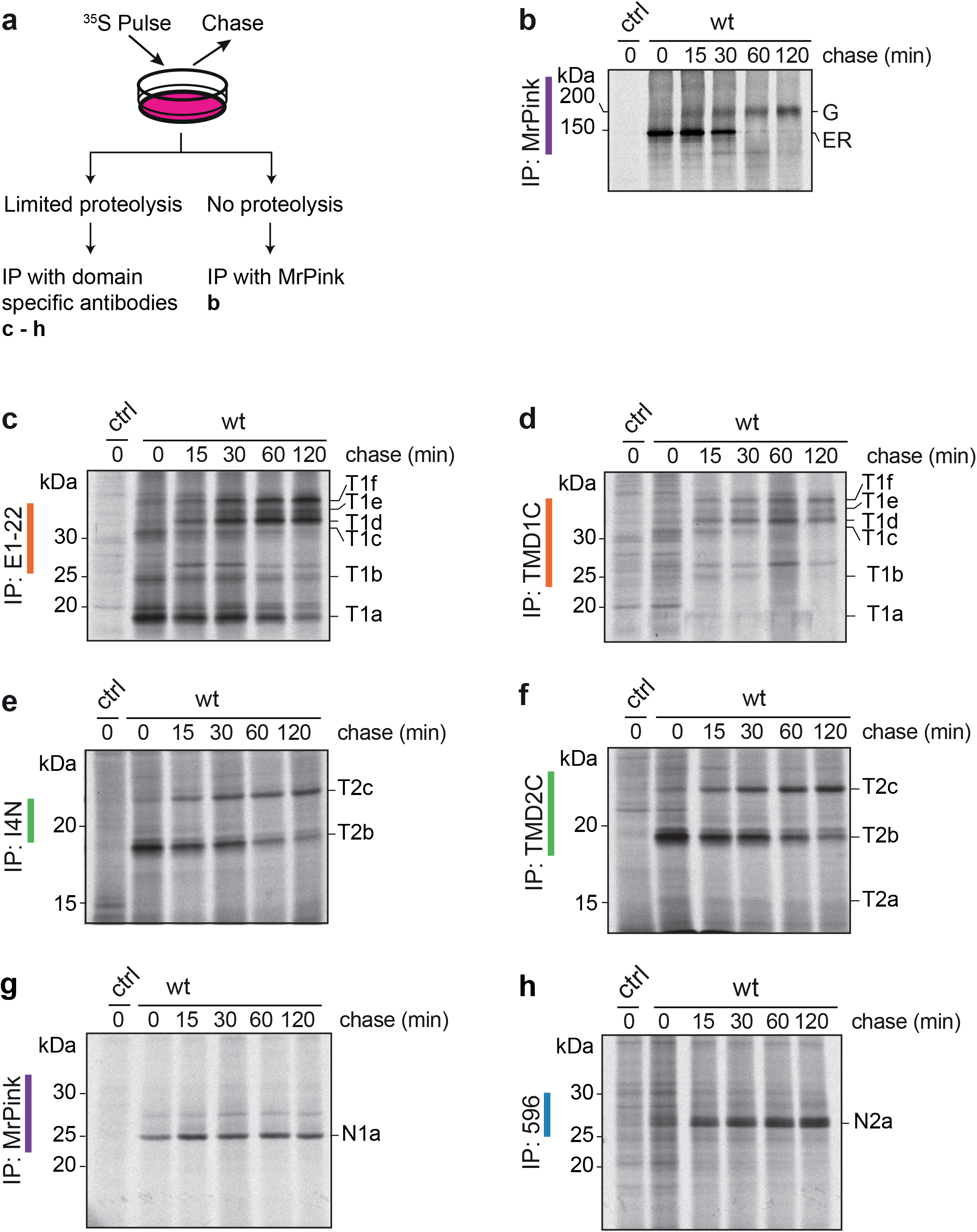
Antibodies against TMDs identify folding intermediates in vivo. **(a)** Workflow of radioactive pulse-chase-limited-proteolysis assay. **(b)** HEK293T cells expressing CFTR were pulse labeled for 15 minutes and chased for the indicated times. CFTR was immunoprecipitated using MrPink and immunoprecipitates were resolved by 7.5% SDS-PAGE. Remaining lysates were subjected to limited proteolysis (LP) with 25 µg/mL proteinase K and protease-resistant fragments were immunoprecipitated with **(c)** E1-22, **(d)** TMD1C, **(e)** I4N, **(f)** TMD2C, **(g)** MrPink, and **(h)** 596 and resolved by 12% SDS-PAGE. wt, wild-type CFTR; T1a-T1f are TMD1-specific protease resistant fragments; T2a-c are TMD2-specific protease resistant fragments; N1a and N2a are protease resistant fragments specific for NBD1 and NBD2, respectively.

As CFTR traveled from the ER to the Golgi complex (Figure 2b), the proteolytic fragments immunoprecipitated by the domain-specific antibodies changed (Figure 2c-h). Especially the TMD1 fragment profiles changed with time, with smaller fragments more prominent immediately after pulse labeling, and larger TMD1 fragments arising from CFTR after the chase. E1-22 and TMD1C immunoprecipitation detected three fragments at 0-h chase, which we named T1a, b, and c. The amounts of T1a-c decreased during the chase and three larger TMD1 fragments emerged, which we named T1d, e, and f (Figure 2c,d).

Like the TMD1 antibodies, TMD2C and I4N immunoprecipitated smaller proteolytic fragments after the pulse as well, and larger products after a chase. Immediately after the pulse, one fragment, which we named T2b, was detected by both antibodies (Figure 2e,f). An additional, smaller fragment, T2a, was faintly recognized by TMD2-C in Figure 2f, and became more prominent when using more lysate (Figure 5a,b). After 2 hours of chase, the T2b early fragment started to disappear with concomitant appearance of the larger fragment T2c (Figure 2e,f). TMD1C, TMD2C and I4N immunoprecipitated substantial amounts of proteolyzed CFTR fragments after 2 hours of chase, in contrast to full-length CFTR. Once fragmented, the epitopes are less shielded and therefore more accessible to the antibodies, implying that these three antibodies are conformation sensitive.

NBD1-specific immunoprecipitates of proteolytic digests contain a major band (N1a) at ∼25 kDa immediately after the pulse, which persists throughout the chase (Figure 2g; [29-31]). For NBD2, digestion yields a single band at ∼25 kDa, N2a, after proteolysis of samples at the end of the labeling period, which increases during the chase (Figure 2h; [29-31]). The time-dependent alterations in fragment patterns suggested that TMD1, TMD2, and NBD2 underwent post-translational conformational changes. The increasing size of the fragments may be due to formation of more compact structures within domains or due to increasing CFTR domain-domain interactions [6, 14].

We therefore set out to determine the identity of each proteolytic fragment. Decoding proteolytic fragments will reveal the protected areas within each domain at different phases of CFTR folding; the identification of protected and deprotected regions reports on the domain folding and assembly mechanisms. To this aim we combined information on the location of the epitopes recognized by the antibodies (Table S1) and on Proteinase-K consensus cleavage sites (Table S2), with secondary structure predictions, and with electrophoretic mobility shifts of fragments digested from CFTR truncations and point mutants.

### The CFTR N-terminus is increasingly protected post-translationally

All fragments from the TMD1-specific immunoprecipitates contained the E1-22 epitope (aa A107-S118) in ECL1, but only the larger T1e and T1f fragments were detected by MM13-4 (aa G27-L34) (Figure 3a, 2-h chase). This showed that the N-terminal boundaries of proteolytic TMD1-derived fragments T1e and T1f must have been upstream of epitope MM13-4, between residues M1 and G27. The N-terminal boundaries of T1a-d must be between epitopes MM13-4 and E1-22, i.e. residues L34 and A107. To zoom in, we generated 19 N-terminally truncated versions of CFTR from the far N-terminus (ΔN2) to the first TM helix (ΔN76). Constructs were expressed in HEK293T cells and analyzed by SDS-PAGE for the impact of truncations on the patterns of proteolytic TMD1 fragments at 0-h, 1-h, or 2-h chase times (Figures 3 and S3). Precise positions of peaks were compared by line scans of gel lanes. This procedure allowed identification of the N-termini of the individual fragments.

**Figure 3.**
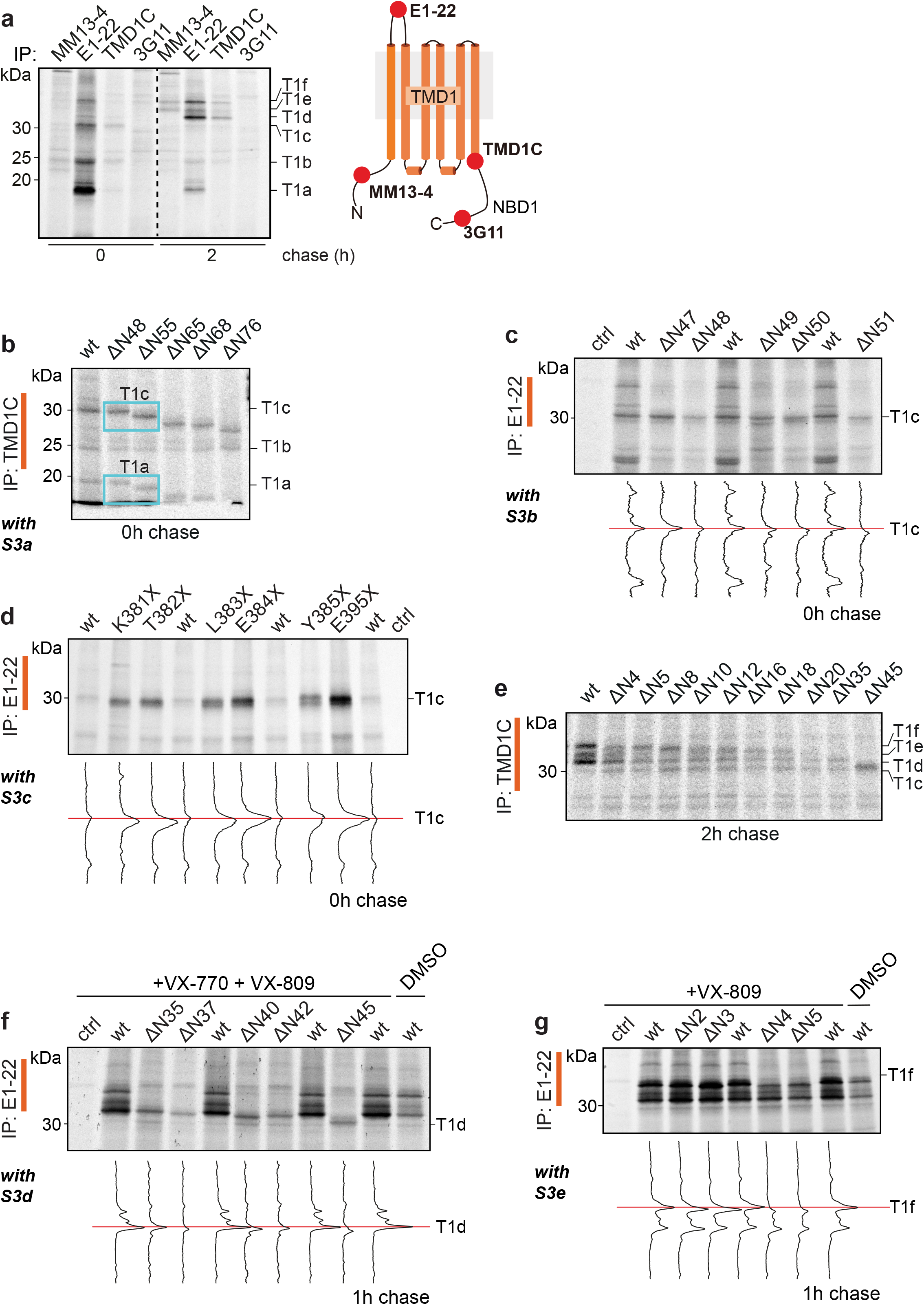
Identification of TMD1 fragments. **(a)** HEK293T cells expressing CFTR were pulse labeled for 15 minutes and lysed immediately or after a chase of 2 hours. Lysates were subjected to limited proteolysis with 25 µg/mL Proteinase K and protease-resistant fragments were immunoprecipitated with indicated antibodies. Position of the antigenic epitopes in TMD1 is marked with red spheres in the cartoon on the right. **(b)** HEK293T cells expressing N-terminally truncated versions ΔN48 to ΔN76 of CFTR were pulse-labeled for 15 minutes and lysed. Lysates were subjected to limited proteolysis with 25 µg/mL Proteinase K and immunoprecipitated with TMD1C. The downward shifts of the TMD1-derived fragments are marked in cyan boxes. **(c)** HEK293T cells expressing N-terminal CFTR truncations ΔN47 to ΔN51 were pulse labeled for 15 minutes and lysed immediately. Detergent lysates were subjected to limited proteolysis with 25 µg/mL proteinase K and immunoprecipitated with E1-22. Lane intensity profiles (ImageQuant analysis) of the fragments of interest are shown below each panel. **(d)** same as **(c)** but now with C-terminal truncations K381X to E395X. **(e)** same as **(b)** but now for ΔN4 to ΔN45 CFTR and after a chase of 2 hours. **(f)** same as **(c)** with N-terminal truncations ΔN35 to ΔN45, but the 15-min pulse labeling was followed by a 1-hour chase. **(g)** same as **(f)** but now with N-terminal truncations ΔN2 to ΔN5. Experiments in panels **(f)** and **(g)** were done in the presence of VX-770 (3 µM) and/or VX-809 (3 µM) as indicated. The undigested samples corresponding to panels b-d and f-g are in Figure **S3a-e**. All samples were resolved by 12% SDS-PAGE. wt, wild-type CFTR.

The size of the T1c and T1a proteolytic fragments decreased when derived from CFTR lacking 55 (ΔN55) or more N-terminal residues, but not yet after truncating 48 residues, ΔN48 (Figure 3b), indicating that the N-termini of T1c and T1a arose from proteolysis between residues N48 and R55. The parallel and gradual decrease of T1a and T1c upon further truncation (Figure 3b) suggested that T1a and T1c share their N-termini. A deletion series of D47 to E51 (Figures 3c and S3a) confirmed Leu49 as the N-terminal cleavage site for T1a [29] and for T1c. To find the C-terminal boundary of T1c, we used C-terminal truncations of CFTR. Introducing a stop codon at positions E384, Y385, or N386 close to the C-terminus of TMD1 (aa M394) did not change T1c mobility, whereas shorter constructs (truncated at K381, T382, or L383) did (Figure 3d). T1c thus represents fragment S50-L383, with Leu49 and Leu383 as the most prominent protease cleavage sites. We had defined fragment T1a as aa S50-S256 [29], which is consistent with T1a lacking the TMD1C epitope (aa S364-K381). Yet, T1a at times does appear in TMD1C immunoprecipitations (Figure 3b), suggesting that despite the proteolytic cleavage at Ser256 in ICL2, the N-terminal and C-terminal halves of TMD1 stay tightly associated upon non-denaturing cell lysis. Cryo-EM structures [7, 14, 16, 17, 21] and structural models [13, 15] confirm that in native CFTR, TMH5 and TMH6 interact with residues in TMH1-4.

T1b must be a mixture of at least two separate, mutually exclusive fragments. T1b appears in immunoprecipitations with MM13-4 (epitope aa G27-L34) and also with TMD1C (epitope aa S364-K381) (Figure 3a,b) and is too small to contain both epitopes. Indeed, the T1b fragment recognized by TMD1C does not change mobility when CFTR is truncated up to 76 residues from the N-terminus (Figure 3b). Because the fragment was not unique, we did not invest in its identification.

To identify the C-terminal boundary of the later-appearing fragments T1d-f, C-terminally truncated constructs cannot be used, as TMD1 requires downstream domains to reach the fully native fold [26] and hence does not acquire sufficient protease resistance to yield T1d-f when truncated. We therefore relied on results of epitope mapping to determine the C-terminus (Figure 3a). T1d-f was immunoprecipitated by TMD1C (epitope S364-K381 at the C-terminus of TMD1), but not by the 3G11 antibody (epitope N396-F405 at the N terminus of NBD1). We concluded that the C-terminal boundaries of T1d-f are in the linker region between TMD1 and NBD1 (Figure 3a). Proteinase-K consensus sites in this region are L383, L387 and M394. The C-terminus of T1c arises from cleavage at Leu383. Because T1c results from digestion of less packed and more open forms of CFTR, the early folding and domain assembly intermediates, Leu383 is the more likely cleavage site compared to M394 and L387. Moreover, M394 is located in a β-sheet [14] and is less likely to be cleaved. We concluded that Leu383 is the most probable cleavage site and last residue of not only T1c but also fragments T1d-f.

To identify the N-termini of fragments T1d-f, we expressed N-terminally truncated versions of CFTR. Proteolytic digestion and immunoprecipitation with the TMD1C antiserum showed that deletion of 4 to 45 residues increased the electrophoretic mobilities of T1f, T1e, and T1d sequentially (Figure 3e), implying that these three fragments differed in their N-termini. Because T1d-f only arise late in biosynthesis, their analysis requires a chase period, during which many truncated constructs were degraded (Figure 3e). To facilitate analysis of fragments T1d-f, we added corrector and potentiator compounds VX-809 and VX-770 [32-36]. VX-809 stabilizes TMD1 and thereby enhances expression of almost all CFTR mutants [32-36]. VX-770 destabilizes T1f at the gain of T1d, improving detection of T1d [24, 37]. The T1d fragment was immunoprecipitated with TMD1C (epitope aa S364-K381) when 35, but not 45 residues had been truncated from the N-terminus, (Figure 3e). The N-terminus of T1d hence lies between aa S35-S45, consistent with the lack of detection by MM13-4 antibody (epitope aa G27-L34) (Figure 3a, right panel). Truncation analysis of aa S35-S45 revealed that deleting 37 residues prevented immunoprecipitation with E1-22 (epitope aa A107-S118) whereas removing 35 residues still allowed detection of T1d (Figure 3f). T1d thus starts from aa D36, with Ser35 being the most prominent protease cleavage site.

The signal of T1e was not strong in wild-type CFTR, and removing residues from the N-terminus eventually caused collapse of T1f onto T1e (Figure 3e). As T1e was immunoprecipitated by MM13-4 (epitope aa G27-L34) (Figure 3a, right panel), we concluded that the N-terminus of T1e lies between residues W19 and K26 in the Lasso helix 1 (Lh1). T1e was present in digests from CFTR lacking the first 16 or 18 amino acids, was only a very fuzzy fragment from the ΔN20 deletion and, as expected, absent upon truncation at amino acid S35 (Figure 3e). The most likely boundary of T1f is residue F17 or S18, from cleavage after Phe16 or Phe17. Yet, the fuzzy nature of the T1e band and its frequent absence –such as from ΔN5 (Figure 3e)– are consistent with the dynamic nature of Lh1 and its embedding in the membrane [14, 15, 38].

T1f had already decreased after truncation of only 4 amino acids of CFTR (ΔN4, Figure 3e). Upon closer inspection even deleting the first two amino acids (ΔN2, Figure 3g) resulted in a downward shift and we conclude that T1f starts from the first amino acid of CFTR.

Results on identification of TMD1 fragments are compiled in Tables 1 and S3 and shown in Figure 4. In summary, T1a starts from N-terminal Lasso helix 2 (Ser50), includes ICL1 and TMH1-4, and ends at Ser256, halfway down the descending ICL2 helix extending from TMH4 [29]. T1c has the same N-terminus as T1a but is protease-protected at the C-terminus of T1a, contains both ICL1 and ICL2, and ends at Leu383 in the linker between TMD1 and NBD1. The difference between the early T1a and T1c fragments and the late T1d-f fragments is the N-terminal region upstream of Lh2, a 50-amino acid stretch that includes Lasso helix 1 and the ultimate N-terminus of CFTR (Figure 4, Table 1). While the protein folds, the N-terminus of CFTR becomes more protected and resistant to protease.

**Table 1.**
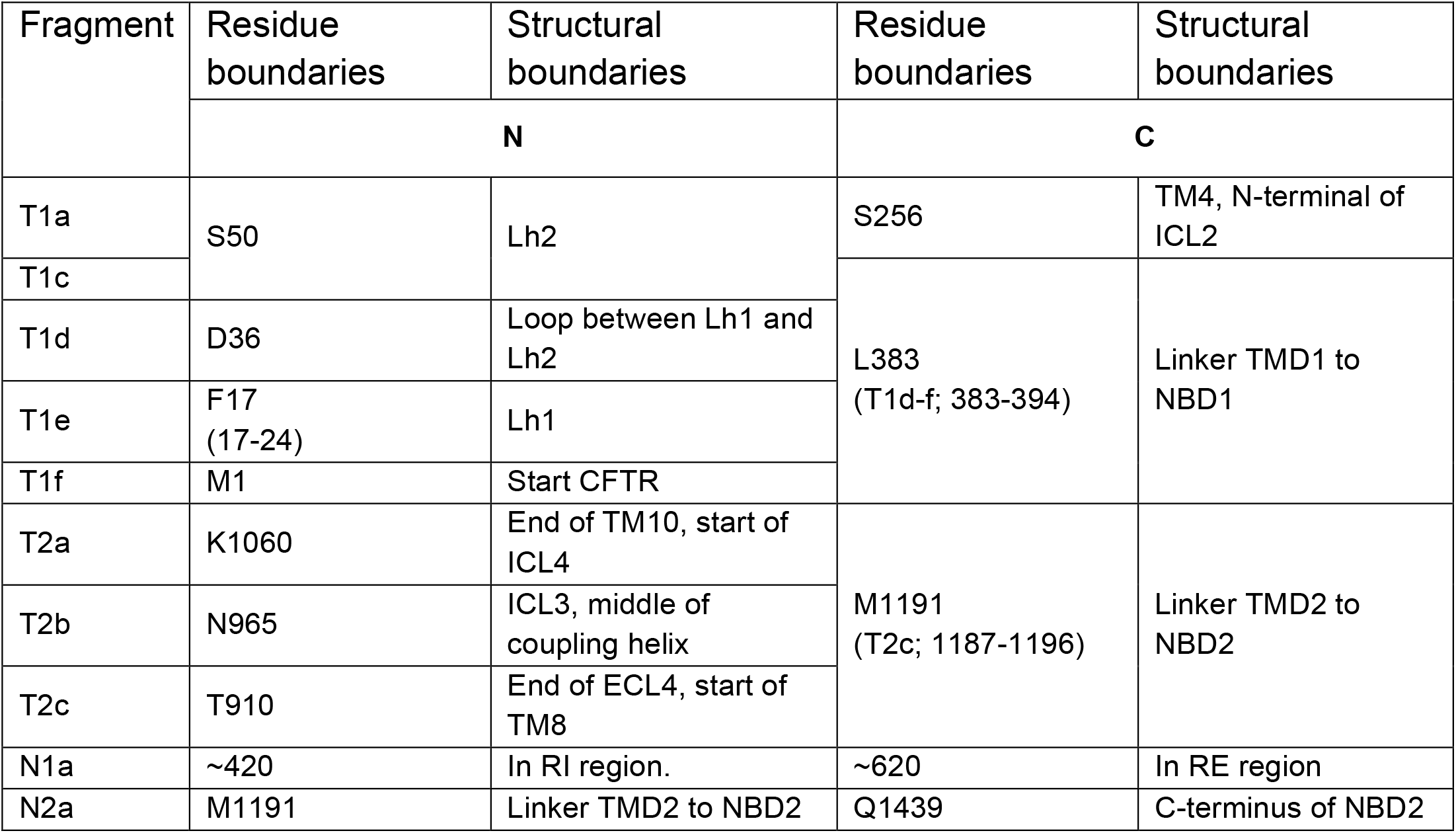
Summary of fragment identities. Summary of residue boundaries of the proteolytic fragments and their locations relative to the structure of CFTR. Where the boundaries are defined by ranges of amino acids, the most likely residue is followed by the range in brackets.

**Figure 4.**
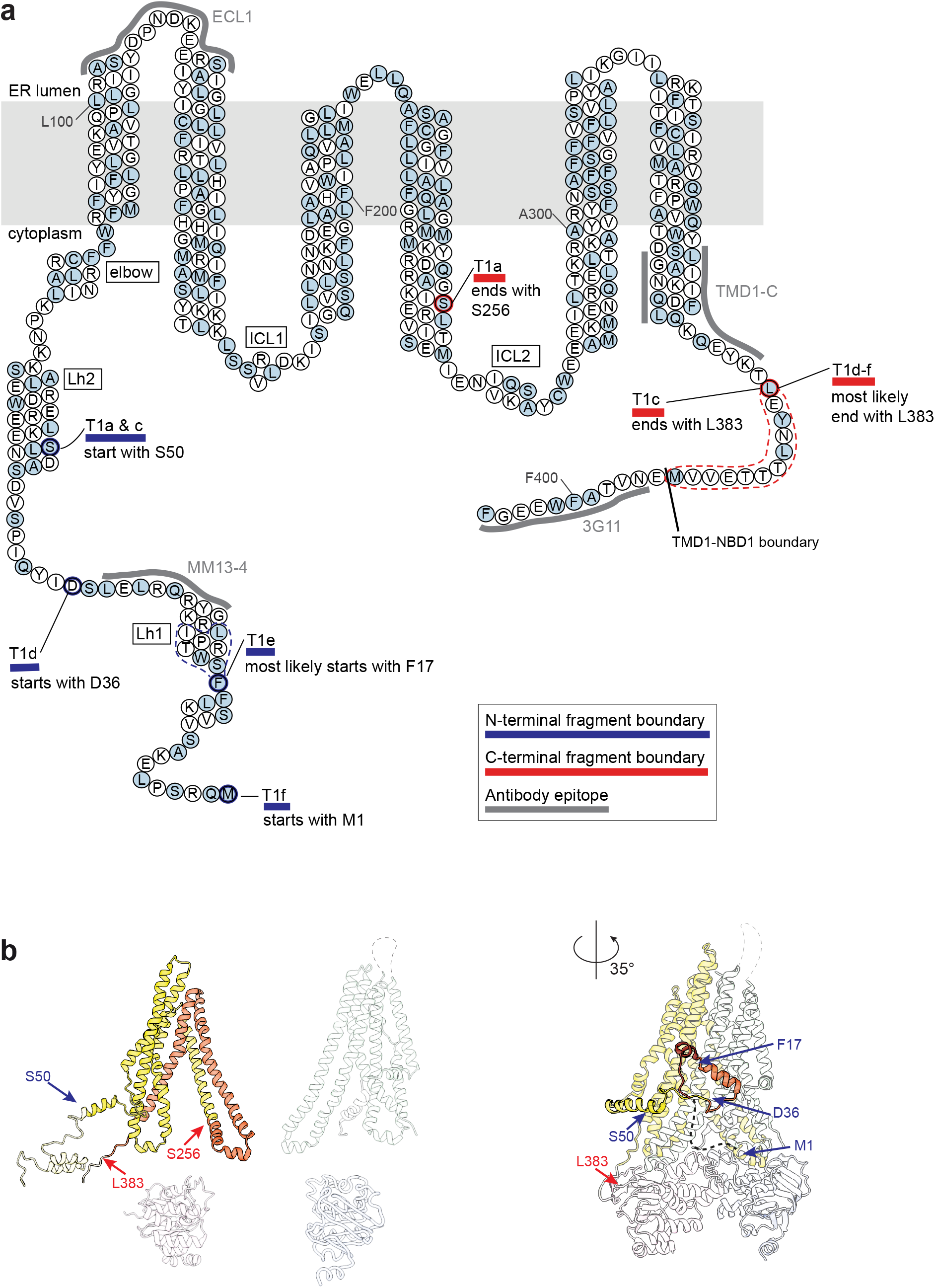
Summary map of TMD1 fragments. **(a)** TMD1 amino acid sequence highlighting the Proteinase-K cleavage sites that result in TMD1 fragments T1a-T1f. All alpha helices are depicted as columns of 3 residues wide, with the exception of ICL1, which includes a coupling helix of only 5 residues (SRVLD). T1a extends from aa S50 to S256, T1c from S50 to L383, T1d from D36 to L383, T1e from F17 to L383, and T1f from M1 to L383. Key: light blue circles: Proteinase-K consensus cleavage residues; grey lines: antigenic epitopes; blue lines: N-terminal boundaries of fragments; red lines: C-terminal boundaries of fragments; dotted lines: possible cleavage area; Lh1: location of lasso helix 1; Lh2, location of lasso helix 2; elbow: location of the N-terminal elbow helix. **(b)** TMD1 proteolytic fragments in structure representation. Left panel shows fragments T1a and T1c with their fragment boundaries. Structure is based on CFTR models with the N-terminus in the cytoplasm [13, 15]. Right panel shows fragments T1d-f with their fragment boundaries. Structure is based on cryo-EM structure (PDB: 5UAK) [14].

**Figure 5.**
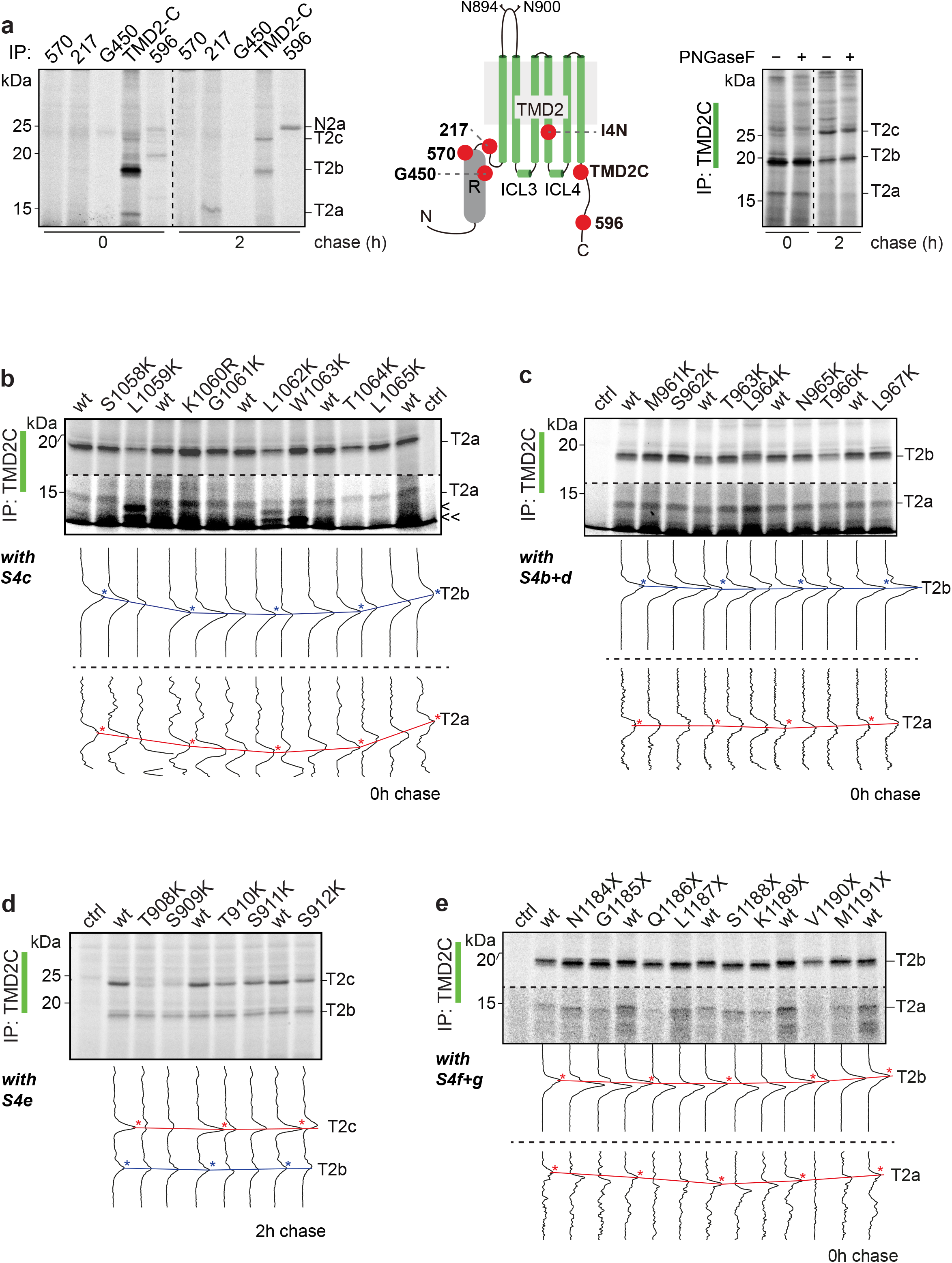
Identification of TMD2 fragments. **(a)** HEK293T cells expressing CFTR were pulse labeled for 15 minutes and lysed immediately or after a chase of 2 hours (left panel). Lysates were subjected to limited proteolysis with 25 µg/mL Proteinase K and protease-resistant fragments were immunoprecipitated with indicated antibodies. Position of the antigenic epitopes in TMD2 is marked with red spheres in the cartoon (middle panel). The right panel shows TMD2-fragment immunoprecipitates treated with PNGaseF. **(b)** HEK293T cells expressing CFTR mutants S1058K to L1065K were pulse labeled for 15 minutes and lysed immediately. **(c)** same as **(b)** but with mutants M961K to L967K. **(d)** same as **(b)** with mutants T908K to S912K, but now after a chase of 2 hours. **(e)** same as **(b)** with C-terminal truncations N1184X to M1191X. Detergent cell lysates subjected to limited proteolysis with 25 µg/mL proteinase K were immunoprecipitated with TMD2C. Samples were resolved by 12% SDS-PAGE. Lane intensity profiles (ImageQuant analysis) of the fragments of interest are shown below each panel. The undigested samples corresponding to panels **(b)-(e)** are in Figure **S4c-f**. Wt, wild-type CFTR; T2a, T2b, and T2c are TMD2-specific protease-resistant fragments.

### ICL4 in TMD2 is protected during post-translational folding

Our strategy to determine the N-terminal boundaries of the TMD2 fragments started with immunoprecipitations using the antibodies 217, 570, and G450 against R (Figure 5a, left and middle panels), which is directly N-terminal of TMD2. None of these brought down TMD2 fragments, showing that the fragments do not include parts of R (Figure 5a, left panel). The fourth extracellular loop (ECL4) in TMD2 contains N-linked glycans at amino-acid residues N894 and N900. We then interrogated TMD2C immunoprecipitates of limited-proteolysis fragments for the presence of N-linked glycans using PNGaseF digestion. In contrast to full-length CFTR (Figure S2), none of proteolytic fragments shifted downwards in the gel following glycanase treatment (Figure 5a, right panel) showing that the TMD2 fragments do not contain residues N894 and N900. Because T2a was immunoprecipitated using TMD2C (epitope E1172-Q1186), but not I4N (epitope Q1035-S1049), we place the N-terminal boundary of T2a after TMH10 (Figures 5a and 2e-f). TMD2 is connected via its N-terminus to the R region and through its C-terminus to NBD2. Generating truncations at the N-or C-termini of TMD2 not only disturbs the interactions with other domains, but also may affect early folding events [26, 27]. For precision mapping the N-terminal boundaries of TMD2 fragments, we therefore used an alternative strategy than deletion analysis. We found fortuitously that changing residues of ICL3 and ICL4 into lysine affected mobilities of proteolytic TMD2 fragments. This observation provided a starting point for investigating the N-terminal boundaries of the proteolytic fragments T2a and T2b using site-directed mutagenesis.

To determine the N-terminal boundary of T2a, we performed a lysine scan on ICL4, mutating one residue at a time to a lysine and subjecting to pulse-chase, limited proteolysis and immunoprecipitation (Figure 5b). The T2a fragment shifted slightly upwards when residues G1061-T1064 were mutated to a lysine, and did not shift in S1058K nor K1060R. A K-to-R mutation is not expected to change mobility, because the charge did not change and SDS binding was likely the same. We concluded that Leu1059 must be the cleavage site that generates T2a, which then starts at K1060. This was underscored by the phenotype of the L1059K mutant: it did not yield T2a but two smaller fragments instead (Figure 5b, < and <<). Similar although not identical patterns emerged from L1062K and W1063K. The small bands appeared at the expense of T2b in the L1059 and L1062 mutants, while radiolabeled full-length protein levels were the same (Figure S4c). These mutations must have changed CFTR conformation and exposed otherwise protected proteolytic sites, likely including L1062 and W1063.

T2b is larger than T2a and was recognized by I4N (epitope aa Q1035-S1049) (Figure 2e). This placed the N-terminal boundary of the fragment upstream of residue Q1035, with ICL3 being the most accessible region for the protease [14]. We therefore mutated residues in ICL3 to lysine, one at a time, and analyzed whether the mutation affected mobility of the T2b fragment (Figure 5c). Patient mutant G970R yielded an upshifted T2b (not shown), which placed the N-terminal boundary of T2b at or N-terminal of G970. We examined lysine mutants of residues L957 to G970 (Figures 5c and S4b, d) and found L964K T2b to shift upwards significantly. The L964K mutation likely eliminated the preferred Proteinase K cleavage site to generate T2b, perhaps to the benefit of S962. T2b electrophoretic mobility appears less sensitive to charge mutations than the other fragments as mutants between L964 and G970 did not cause a mobility shift. We conclude that T2b starts at Asn965 upon Proteinase-K cleavage after Leu964.

As T2c did not contain glycans (Figure 5a) and had a higher mass than T2b, we hypothesized that the N-terminal boundary was in ECL4 downstream of the N-linked glycosylation sites. Residues C-terminal to the N-glycosylation site N900 in ECL4 were mutated to lysine to identify the N-terminal boundary of T2c (Figure 5d, S4a, e). Mutants N901K to V905K yielded T2c with unchanged mobility (Figure S4a), which eliminated S902 and A904 as possible T2c protease cleavage sites and suggested that these residues were not part of T2c. In contrast, lysine mutations in residues S911 and S912 did decrease mobility, implying that they were included in T2c. In mutants T908K and S909K between the non-affected (N901-V905) and affected residues (S911-S912), the intensity of T2c was decreased significantly. As threonines are not favored Proteinase-K cleavage sites, we conclude that Ser909 is the most likely cleavage site to generate T2c, positioning its N-terminal boundary at Thr910.

By analyzing the presence of the epitopes on the fragments, we conclude that the C-terminal boundaries for the TMD2 fragments were between aa K1189-I1203, as they all were immunoprecipitated by TMD2C (epitope E1172-Q1186) but not 596 (epitope W1204-T1211) (Figure 5a, left and middle panels). Replacing V1190 with a stop codon still generated T2b, whereas shorter constructs led to a downward shift (Figure 5e, S4f,g). The shift changes were also seen for T2a, which would suggest that the last residue of both fragments would be K1189. This is not likely as K is not cleaved by Proteinase K. More probable is M1191 as preferred cleavage site and C-terminal residue of the TMD2 fragments, because it is the first cleavable residue upstream of K1189. The aberrant mobilities of the fragments derived from M1191X and V1190X may well be caused by the positive charge of the lysine residue close to their C-terminus. Removal of the K in K1189X leads to an immediate shift down in the gel. Indeed, if anything, V1190X ‘T2b’ may run even slightly higher than wild-type T2b.

We therefore concluded that T2a consists of K1060-M1191, while T2b is N965-M1191. The C-terminus of T2c could not be determined this way as it was not detected in C-terminally truncated CFTR after the chase (data not shown). We nonetheless concluded, considering the epitopes, that T2c most likely is cleaved at M1191 as well and encompasses T910-M1191 (Figure 6).

**Figure 6.**
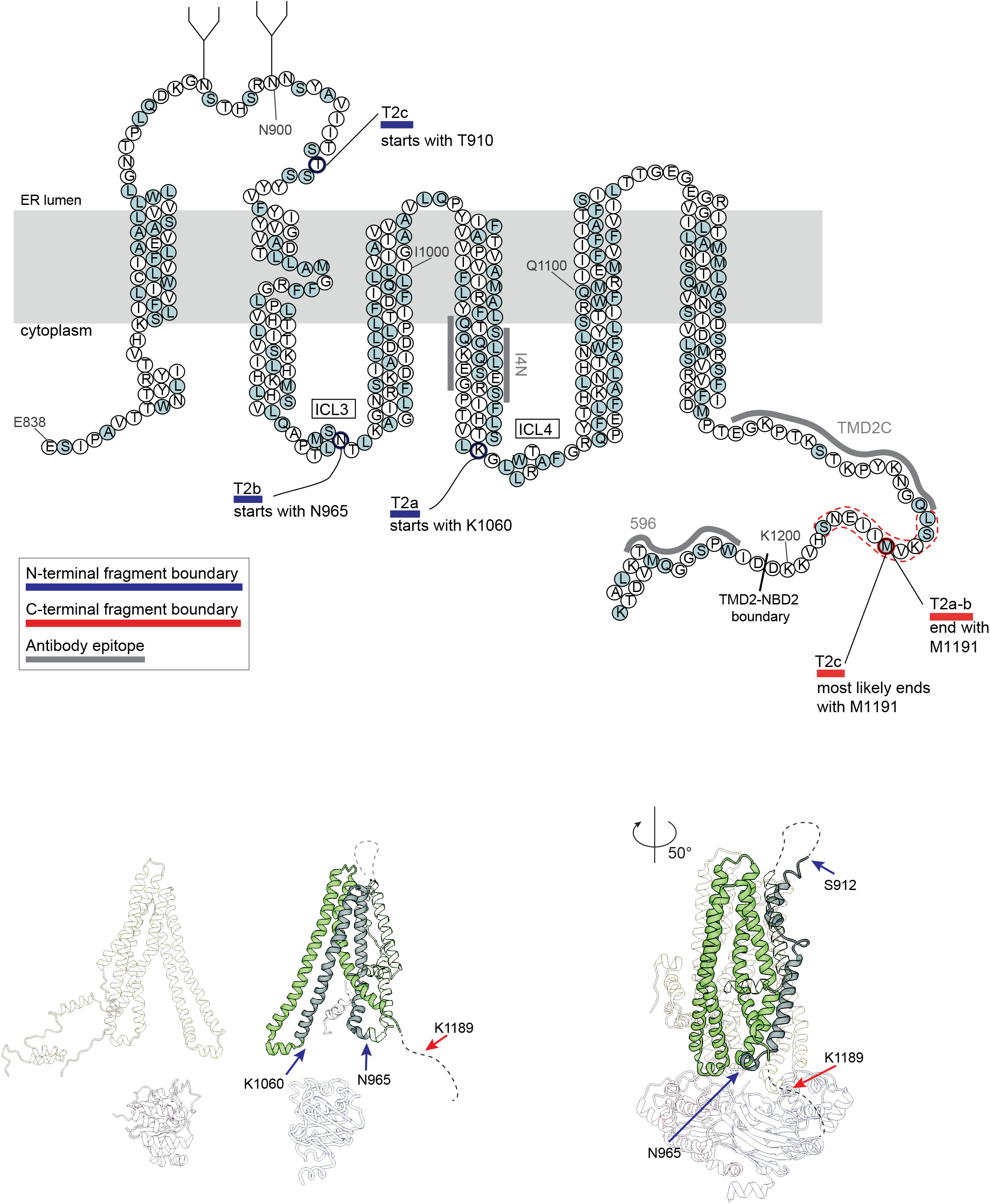
Summary map of TMD2 fragments. **(a)** Representation of the TMD2 amino-acid sequence highlighting the Proteinase-K cleavage sites that result in TMD2 fragments T2a-T2c. All alpha helices are depicted as columns of 3 residues wide. T2a extends from aa K1060 to M1191, T2b from N965 to M1191 and T2c from T910 to M1191. Key: light blue circles: Proteinase-K consensus cleavage residues; grey lines: antigenic epitopes; blue lines: N-terminal boundaries of fragments; red lines: C-terminal boundaries of fragments; dotted lines: possible cleavage area. **(b)** TMD2 proteolytic fragments in structure representation. Left panel shows fragments T2a and T2b with their fragment boundaries. Structure is based on CFTR models with the N-terminus in the cytoplasm [13, 15]. Right panel shows fragment T2c with its fragment boundaries. Structure is based on cryo-EM structure (PDB: 5UAK) [14].

Results on identification of TMD2 fragments are compiled in Table S4. In summary, T2a starts near the beginning of ICL4 until the end of TMD2. T2b shares the same C-terminal end as T2a but starts within ICL3. With time, we start to see the appearance of T2c, which starts at the C-terminal end of the fourth extracellular loop (Figure 6, Table 1). The change from T2a-b to T2b-c shows that ICL4 becomes protected while CFTR folds.

### N1a and N2a contain almost entire NBDs

The nucleotide-binding domains yielded one ∼25-kDa major protease-resistant fragment each, which we named N1a and N2a (Figure 2g, h). N1a contains the almost complete NBD1, without the N-terminal strand and C-terminal helices (manuscript in preparation). The proteolytic cleavages occur in the relatively unstructured regions RI and RE of NBD1 (Figure 7a).

**Figure 7.**
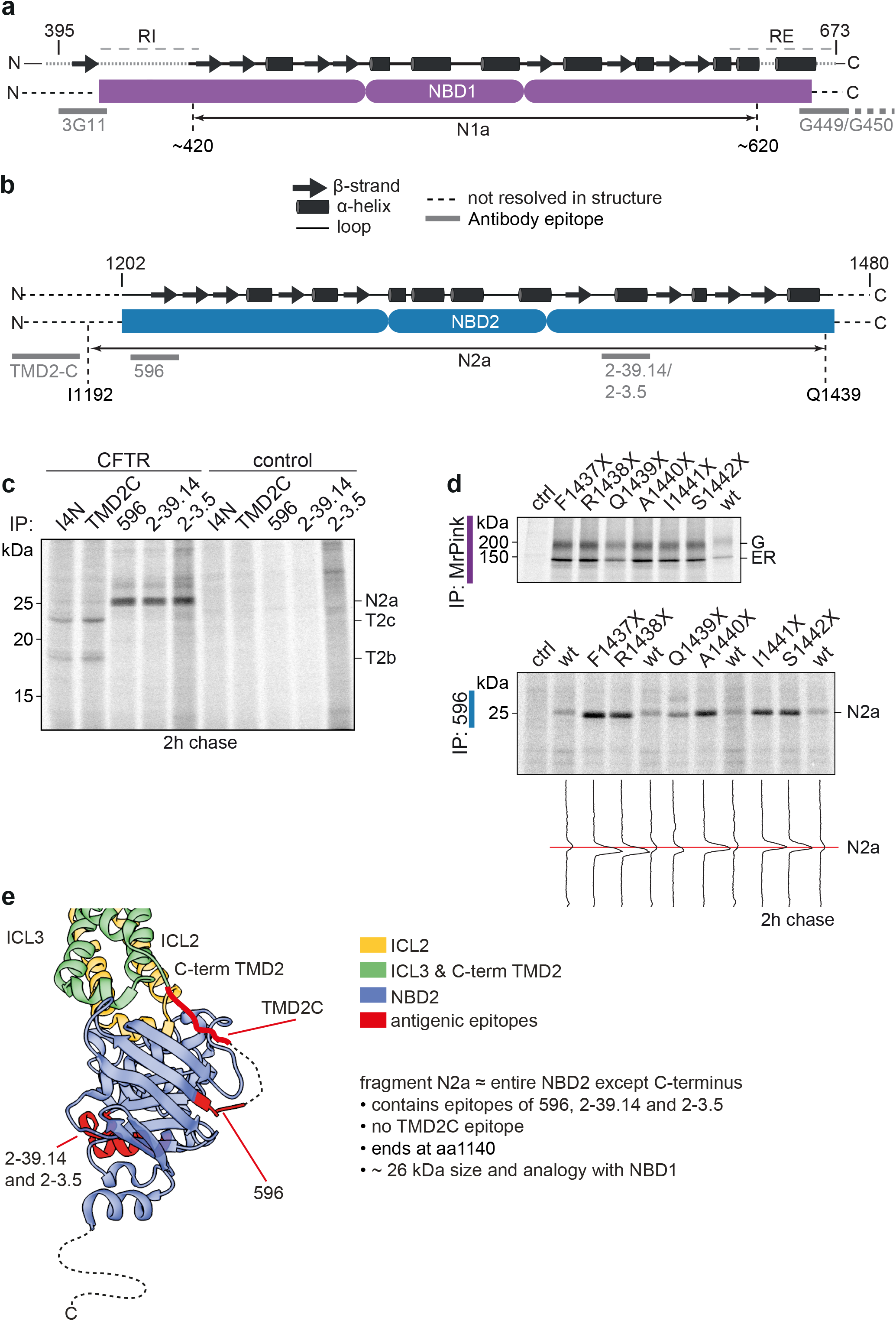
The late NBD2 fragment contains almost all of NBD2. **(a)** Secondary structure of NBD1 with flanking Regulatory Insertion (RI) and Regulatory Extension (RE), with antigenic epitopes indicated and showing the identity of the N1a fragment, residues ∼420 to ∼620. **(b)** Secondary structure of NBD2 with antigenic epitopes indicated and showing the identity of the N2a fragment, residues I1192-Q1439. **(c)** HEK293T cells expressing CFTR or not (control) were pulse labeled for 10 minutes and chased for 2 hours. After digestion with 25 µg/mL Proteinase K, proteolytic fragments were immunoprecipitated in parallel with I4N, TMD2C, 596, 2-39.14, and 2.3.5 antibodies and resolved by 12% SDS-PAGE. **(d)** HEK293T cells expressing C-terminally truncated CFTR constructs F1437X to S1442X were pulse-labeled for 15 minutes and chased for 1 hour and subsequently digested or not with 25 µg/mL Proteinase K for 15 minutes on ice. Non-digested lysates (upper panel) were immunoprecipitated with MrPink and proteolyzed samples with NBD2-specific antibody 596 (lower panel) and resolved by 7.5% or 12% SDS-PAGE, respectively. Lane intensity profiles (ImageQuant analysis) of N2a are shown below the panel. **(e)** Structure of NBD2 and interacting TMD elements ICL2 and ICL3 (PDB: 5UAK) (Liu et al., 2017) indicating the antigenic epitopes in red and highlighting the NBD2 fragment N2a in black outline. N2a contains almost the entire NBD2 domain (residues D1202-L1480).

To determine the N-terminus of N2a, we analyzed the presence of antigenic epitopes on the fragment (Figure 7b,c). As shown in Figure 7c, N2a was immunoprecipitated by 596 (epitope aa W1204-T1211), 2-39.14 (epitope aa E1371-R1385) and 2-3.5 (epitope aa V1379-T1387) but not by TMD2C (epitope aa E1172-Q1186). We therefore conclude that the N-terminus of N2a was in the linker region of TMD2 and NBD2 (aa K1189-I1203).). We conclude that M1191, which is the C-terminal cleavage site of all three TMD2 fragments, is the logical N-terminal boundary of N2a, because it is in a cluster of consensus sites. W1204 is blocked by the proline in 1205 and we cannot exclude that in (near-)native CFTR M1191 gets protected and the cleavage is at S1196. The C-terminal region of NBD2 (F1437-L1480) is not resolved in the CryoEM structure [14], and is thought to be disordered. Using the C-terminal-truncation strategy, we found that replacing A1440 with a stop codon still immunoprecipitated N2a, whereas shorter constructs led to a downward shift (Figure 7d, bottom panel). N2a hence encompasses complete NBD2 except the C-terminus, from the linker between TMD2 and NBD2 to residue A1440 (Figure 7e, Table S5).

### Mutations in the diacidic ER export motif cause a global folding defect in CFTR

Having characterized the antibodies and the biochemical folding process, we set out to investigate the effect of mutations in the diacidic (DxD) motif in NBD1 (D565-D567; Figure 8a) motif of NBD1 on the folding of CFTR. Substitution of Asp567 to Ala impairs sec24 binding and COPII dependent export of CFTR from the ER [39, 40]. It is not clear however whether these phenotypes are a direct consequence of the mutation in the interaction motif or rather are secondary and reflect a primary defect in folding of NBD1 and assembly of full-length CFTR. Such a difference is suggested by the different (chaperone) interactome of the D565A-D567A mutant compared to that of wt CFTR [41]. We created alanine mutations for each of the aspartate residues as well as the D565G patient mutation [42] and determined the effects on CFTR folding using the combined tools and information in a radioactive pulse chase-limited proteolysis-immunoprecipitation assay.

**Figure 8:**
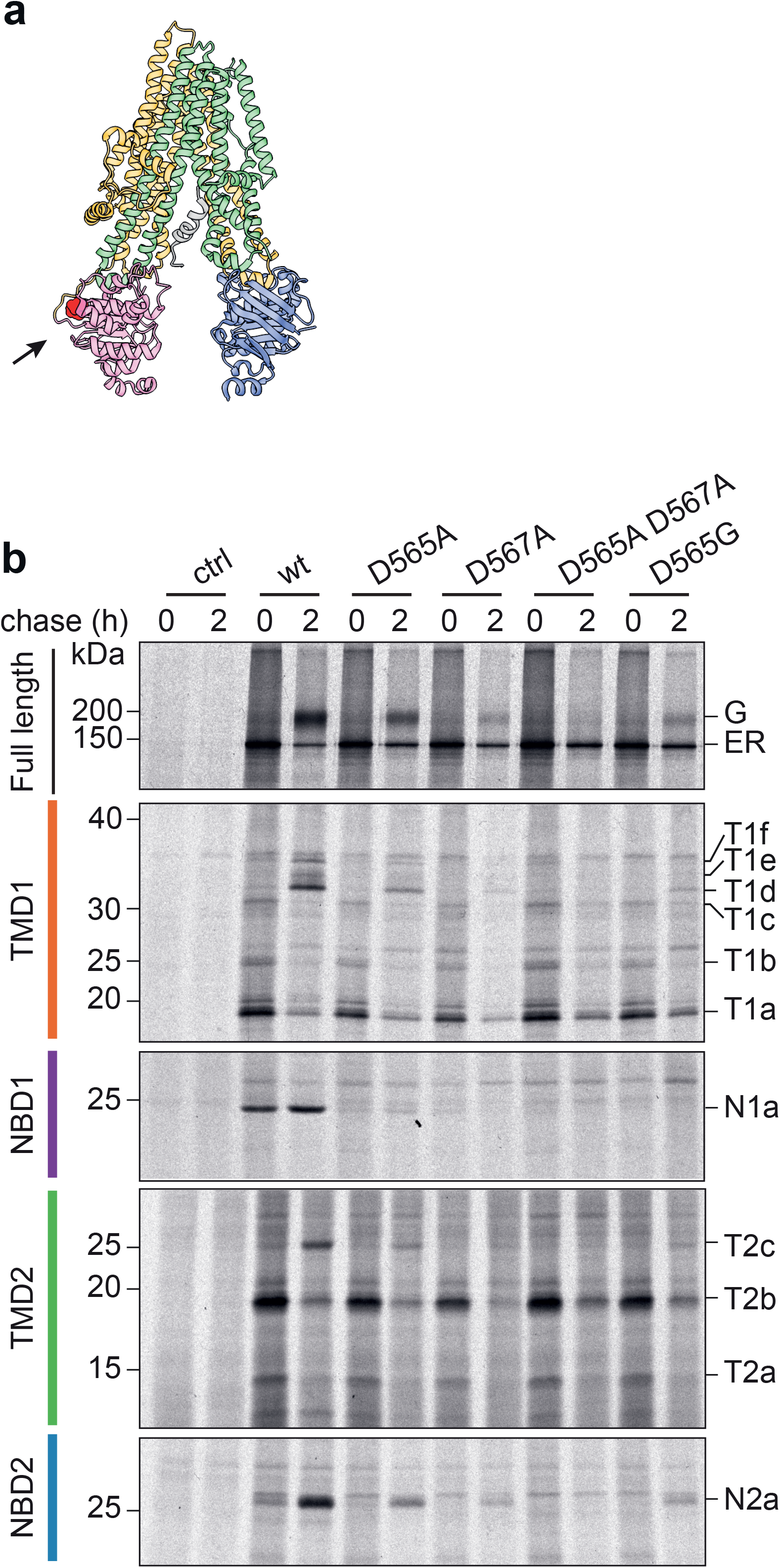
Folding analysis of CFTR mutated in an NBD1 di-acidic ER-export motif. **(a)** Structure of NBD1 showing the location of the di-acidic ER exit motif (arrow & red-shaded area; PDB: 5UAK) [14]. **(b)** HEK293T cells expressing CFTR variants were pulse-labeled for 15 minutes and lysed immediately or after a chase of 2 hours. CFTR was immunoprecipitated from non-proteolyzed lysate with MrPink and resolved by 7.5% SDS-PAGE. Proteolysis was done with 25 µg/mL Proteinase K and domain-specific fragments were immunoprecipitated with E1-22 (TMD1), MrPink (NBD1), TMD2C (TMD2) and 596 (NBD2) and analyzed by 12% SDS-PAGE.

As shown in Figure 8b (top panel), the acquisition of the complex glycosylated form was impaired by mutations in D565 and D567. The effect of the D567 mutants was stronger than of D565 even though a change in the latter is associated with CF [43, 44]. Of note for each of the mutants is that the early steps in the folding path of NBD1 were impaired already because the N1a fragment was barely detectable after the pulse period. In accordance with the notion that NBD1 folding is a limiting step in CFTR biogenesis [18, 45, 46] we found deleterious effects on the amounts of late TMD1, TMD2, and NBD2 fragments at the 2-h chase time, albeit slightly milder for the D565A mutant. Application of the new antibodies raised against the TMDs show that mutations in the diacidic export motif cause misfolding of CFTR into an F508del-like phenotype: misfolding of NBD1 and as a result no assembly with the other domains. Investigations into the interactions of CFTR with the COPII sorting machinery (and other relevant interactors) using mutational approaches thus need to be complemented with careful analysis of the folding status of these mutants.

## DISCUSSION

The folding of multi-domain, polytopic membrane proteins such as ABC-transporters is complex. In this study, we raised antibodies to the transmembrane domains of CFTR, which recognized their epitopes in full-length CFTR and in proteolytic fragments arising from limited digestion with Proteinase K. Using these antibodies together with previously developed antibodies against the nucleotide-binding domains, we identified domain-specific proteolytic fragments and present the 2-stage folding pathway of CFTR in live cells (Figure 9). During translation, TMD1, NBD1, and TMD2 acquire a native-like fold as individual domains. After synthesis, stage 2 shows simultaneously increased protease resistance due to assembly of TMD1, TMD2, and NBD2 onto NBD1, as well as the N-terminus that is wrapped around the TMDs in the native structure [14, 47, 48]. Stage 2 coincides with transport of CFTR from the ER to the Golgi complex and the appearance of the CFTR complex-glycosylated Golgi form.

**Figure 9.**
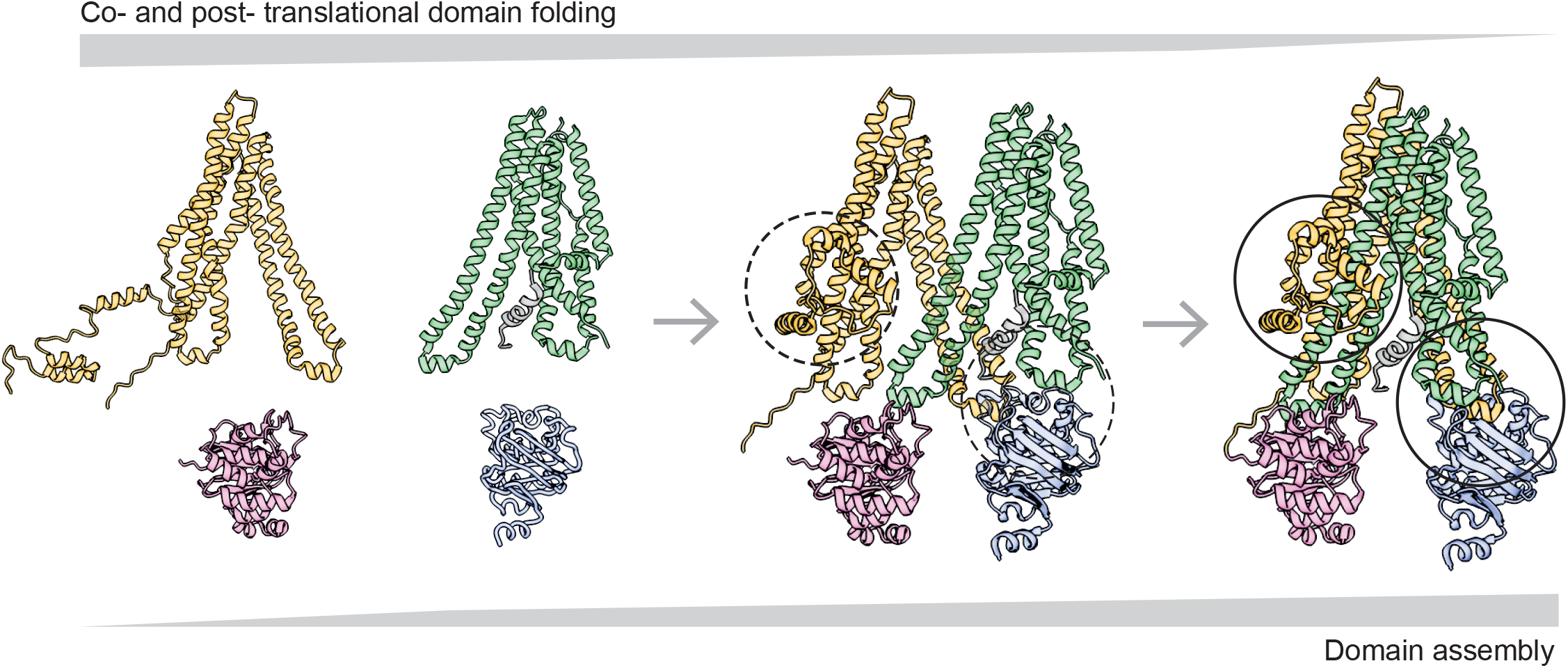
Two-stage folding process of CFTR. The left structure illustrates stage 1 of folding: TMD1, NBD1, and TMD2 fold already co-translationally and acquire a structure as if expressed on their own [7, 29]. ICL1 docks onto NBD1 already during synthesis [29], whereas the N-terminus of TMD1 cannot have native structure yet and may hang off the ER membrane [13] and/or associate with NBD1 [15]. NBD1 reaches its native protease-resistant fold completely co-translationally, which does not change upon domain assembly. Structure is based on CFTR models with the N-terminus in the cytoplasm [13, 15]. The middle structure shows a putative domain-assembly intermediate, in which the domain interfaces form fast and cooperatively, yielding the right structure. The fully domain-assembled CFTR (right structure) has ICL1 of TMD1 and ICL4 of TMD2 docked onto NBD1. The TMDs assemble, such that ICL2 of TMD1 and ICL3 of TMD3 associate with and stabilize NBD2. The circles show the sites that gain significant protease resistance around the time CFTR leaves the ER, reaches the Golgi complex and obtains complex glycans. ICL2, ICL3, and NBD2 acquire increased protease resistance simultaneously (right circle), as does the N-terminus of TMD1, which becomes protease resistant by wrapping around TMD1 and TMD2 (left circle). Structure in middle and right panels are based on cryo-EM structure (PDB: 5UAK) [14].

Previous studies have shown that CFTR folds its domains largely co-translationally; insertion of the transmembrane helices occurs during translation [18] and the individual domains form protease-resistant structures with exception of NBD2 [7, 23, 49]. The structures [14, 16, 17] and structural models [13, 15] of CFTR and several biochemical studies have shown that the transmembrane domains interact with the nucleotide-binding domains through the intracellular loops that connect the transmembrane helices [50-53]. These interactions have been postulated to take place post-translationally [26, 51]. A current model suggests that after co-translational folding of individual domains, a TMD1-NBD1-R-TMD2 structure is produced, with NBD2 incorporation as a final step that is not required for cellular trafficking of CFTR [27]. Our kinetic assay allows a more detailed analysis of the order of events that occur, and we propose the following two-step model for folding (Figure 9).

In stage 1, the first transmembrane domain of CFTR folds co-translationally, with the N-and C-termini packing with each other and with ICL1 [29]. NBD1 reaches a protease-resistant fold during translation as well [7, 23], and its N-terminal region then already interacts with TMD1 through ICL1 [29]. At this time, the N-terminus of TMD1 may well interact with NBD1 as suggested by a structural model [15] and be held by chaperones. TMD2 reaches a protease-resistant fold during translation, yet requires TMD1 and NBD1 to acquire native stability. In stage 2, ICL4 docks onto NBD1 and associates with ICL1. TMD1 and TMD2 assemble with NBD2 via ICL2 and ICL3, and all three acquire their native protease-resistant fold. The TMD1 N-terminus moves to its native position in the structure, wrapping around both transmembrane domains.

### Stage 1: co-translational folding of domains

The individual domains of CFTR fold to a large extent co-translationally [7]. The folding of the two transmembrane domains of CFTR begins with co-translational insertion of the transmembrane helices into the membrane [18], and in this study we show that their packing increases both co- and post-translationally. Immediately upon radiolabeling, CFTR is digested by Proteinase K into fragments that also arise upon digestion of individual TMDs: fragments T1a and T1c from TMD1, and T2b from TMD2. The larger T1c comprises almost the entire TMD1, sans N-terminus, suggesting that all of the six transmembrane helices are packing with each other, protecting the two intracellular loops. In-vivo studies using several membrane proteins in *E. coli* have shown that specific ‘packing’ interactions between the N- and C-terminal TMHs occur during translation, and that these are essential in forming a stable tertiary structure [8]. This also applies to the CFTR TMDs, with the difference that the long intracellular loops emanating from the TM helices start packing in the cytoplasm rather than in the lipid bilayer. Increased protection of the cytosolic N-terminus of TMD1 against digestion requires the presence of its cytosolic C-terminus [8]. In the closed cryoEM structure [14], TMH6 is wedged between TMH1-3 and TMH4, separating TMH4 from TMH1-3; this may well explain the accessibility of Proteinase K to Ser256 in ICL2 and the formation of T1a, which in essence is a truncated T1c. When digested with more protease, the T1a fragment persisted, suggesting that TMH1-3 forms a stable trimeric helical bundle, an evolutionarily conserved folding unit found in multi-spanning membrane proteins [29, 54]. For CFTR, early formation of a stable TMD1 structure is important, as underscored by the mode of action of class-I correctors: VX-809 (Lumacaftor) and VX-661 (Tezacaftor) bind and stabilize TMD1 and thereby rescue mutant CFTR [29, 33, 35, 55]. TMD1 may well be required as template for assembly with the other domains and generation of a functional chloride-channel.

Once NBD1 is translated, it folds quickly and starts co-translational domain assembly with TMD1 [7, 29]. The NBD1-specific immunoprecipitate contains the N1a fragment we had previously identified, containing NBD1 lacking its N- and C-termini [23].

Protease resistance of NBD1 notably preceded that of the other domains, showing that NBD1 folds and obtains its native protease resistance independent of the other domains and thus faster. The rapid and independent folding of NBD1 is underscored by the many studies on purified NBD1 [53, 56, 57], and highlights its importance as a docking scaffold for ICL1 and ICL4 of the transmembrane domains. The N-terminal subdomain of NBD1 (residues G404-L436) improves TMD1 folding co-translationally, by improving packing of ICL1 with the N- and C-termini of TMD1 [29]. This early docking of ICL1 onto NBD1 is complemented with the later docking of ICL4 onto NBD1 around F508. This explains the multiple defects F508del causes: not only NBD1 misfolding but also disruption of an essential domain-assembly step, the interaction of ICL4 (in TMD2) with the F508 region in NBD1 [23, 27, 45, 58-61].

Although size and amino-acid composition of TMD1 and TMD2 are similar, their folding, structures, and stabilities in the cell differ. In contrast to TMD1, TMD2 expressed as a single domain in cells is vulnerable to degradation. In the context of full-length CFTR or when attached to only NBD2 [55] or tagged in ECL4 (our unpublished observations), TMD2 gains stability. This suggests that co-translational folding of TMD2 still yields a vulnerable structure that derives stability from other domains of CFTR. The tagging in ECL4 likely stabilizes TMH8, which is a kinked helix and is notoriously aggregation-prone [14, 16, 62]. The attachment to NBD2 may stabilize TMD2 by assembly through ICL3 and/or limiting flexibility of the TMD2 C-terminus (ref). The hydrophilic aspartate at position 924 in TMH8 jeopardizes stable insertion of TMD2 into the lipid bilayer [4] but is stabilized by R347 in TMH6 of TMD1 of domain-assembled, native CFTR [14, 16]. R347P is CF-causing, whereas D924N occurs but has unknown significance (CFTR2 database).

The early folding fragment T2b starts from the coupling helix of ICL3 to the C-terminal end of TMD2, with ICL4 being protected. The four helices comprising T2b, which are TMH9-12, together with TMH1-2 of TMD1, form the domain swap-structure seen in CFTR [14]. The N-terminal boundary of the smaller T2a fragment is near the end of the cytosolic extension of TMH10, just before the ICL4 coupling helix begins, indicating that ICL4 is at least partially exposed. This exposure may occur as ICL4 is not yet docked onto NBD1. Once ICL4 is docked onto NBD1 via the coupling helix, it starts to pack against ICL1 of TMD1.

The CFTR domains, especially the TMDs, must maintain a certain level of flexibility for interacting with the NBDs and opening and closing the channel, which is consistent with the linkers between all domains remaining exposed and cleavable throughout both stage 1 and stage 2 of the CFTR folding pathway.

### Stage 2: post-translational domain assembly

At later chase times, in stage 2 of CFTR folding, the appearance of fragments T1d-f shows protection of both ICL2 (decrease of T1a) and the N-terminus (T1d-f).

Increasing protease resistance of the N-terminus is consistent with the N-terminus wrapping around the transmembrane helices of TMD1 and TMD2 [14]. TMD1, TMD2, and NBD2 simultaneously show conformational changes, with the protection of ICL2 and ICL3, and protease-resistance of NBD2. Once TMD2 starts to pack with TMD1 and NBD2, ICL2 in TMD1 and ICL3 in TMD2 start packing with each other and with NBD2, which results in simultaneous ICL2 and ICL3 protection. This gives rise to the T2c fragment, which consists of TMH8-12, almost the entire TMD2. TMH7 is not in T2c, whereas it is a stable TMH that acts as signal peptide for TMD2 [21, 63]. We hypothesize that it does associate tightly with TMH8-12 but is separated in SDS-PAGE by cleavage in ECL4, caused by the unstable TMH8 [62]. ICL2 and ICL3 may interact before binding to NBD2. A previous study showed that CFTR without NBD2, with ICL2 and ICL3 being exposed to the cytoplasm, exits the ER [27]. For CFTR to be released from chaperones, pass quality control and exit the ER, hydrophobic amino acids need to be buried inside the structure, and this can be accomplished by ICL2-ICL3 lateral packing. Even in the presence of NBD2, however, ICL2 and ICL3 remain somewhat exposed to protease after their assembly, as evidenced by some persistence of T2b and T1a, which arise from cleavages in ICL3 and ICL2, respectively. This suggests that the interaction interfaces between NBD2, ICL2, and ICL3 are weaker than those of ICL1, ICL4, and NBD1. The cryo-EM structures suggest a larger (and hence stronger) interface of ICL2&3 onto NBD2 than of ICL1&4 onto NBD1 [47]. This hints at the lateral packing of ICL1 with ICL4 perhaps being stronger than that of ICL2 with ICL3.

NBD2 and NBD1 are very similar in structure, and one would expect that they follow the same folding pathways. If NBD2 can fold independently of other domains, it would be puzzling why assembly of NBD2 with the transmembrane domains –and the acquisition of a protease-resistant fold– occurs at a later stage in folding, starting at 30 minutes of chase and continuing up to 2 hours. From our data, NBD2 folds with slower kinetics than NBD1, and we conclude that NBD2 requires the other domains to fold. Assembly of NBD2 with the two TMDs may require prior assembly of TMD1 and TMD2, with each other and with NBD1. The interfaces between the transmembrane helices of TMD1 and TMD2 are complex and may require TMH reorganization and membrane lipid displacement, making it consequently a lengthy process.

Incorporation of the new TMD antibodies into the coupled pulse chase-limited proteolysis assay also allowed us to look at higher resolution into consequences of mutations in CFTR on conformations of the maturing protein. For instance the DAA mutant of the diacidic ER export signal accumulates in the ER and fails to be recognized by the COPII machinery. Initially it was thought that this mutation encodes a relatively pure sorting mutant as opposed to a conformational mutant such as F508del [40], which thus should be primarily defective at the level of sec24 binding.

More recent findings however show enhanced interaction of an ER export mutant with GET4 [41], which together with BAG6 and UBL4 promotes degradation of misfolded ER proteins [64]. Moreover, a patient mutant within (D565G) or close to the diacidic ER export motif within NBD1 is misfolded as shown by us and others [46]. In accord with this central role of NBD1 in the assembly of CFTR domains, we now show that mutants of the DAD motif fail to fold NBD1 and as a result fail to assemble their domains correctly, suggesting that conformational changes prevent entry of CFTR into the COPII ER-export pathway, rather than the mere absence of an export signal.

Collectively, we show that a repertoire of domain-specific antibodies together with a coupled pulse chase-limited proteolysis immunoprecipitation assay constitutes a powerful method to uncover folding pathways in vivo in a temporal manner and to investigate folding-function relationships of (mutant) proteins.

We provide experimental evidence from intact cells for a fundamental principle in protein folding shown before by in-vitro refolding studies: that low-contact-order interactions are formed before high-contact-order folding. The pathway of domain folding before domain assembly that CFTR follows makes sense from a point of view of fidelity of folding, order of folding, and it fits to in-vitro folding principles and models such as the hydrophobic-collapse and the nucleation-condensation models of folding. Considering the high structure conservation of the features that are dominant in CFTR folding, we anticipate these folding events to be highly similar for other ABC-transporters and likely for other multispanning membrane proteins as well.

## MATERIALS AND METHODS

### Antibodies

Epitopes of the anti-CFTR antibodies that were used in this study are listed in Table 1. Generation of the polyclonal rabbit antiserum Mr Pink was described before [23]. Polyclonal antibody ECL1 (31/32), directed against the cyclic peptide of the CFTR extracellular loop 1 [65] was generously donated by Dr. Raymond Frizzell (University of Pittsburgh, USA). Rabbit polyclonal G449 (bleed G450), kindly provided by Drs. Angus Nairn (Yale University, USA) and Hugo de Jonge (Erasmus Medical Center, Rotterdam, The Netherlands), was generated against human CFTR residues F653-K716 [66]. Mouse monoclonals 570, 217 and 596, kindly provided by Dr. John Riordan (University of North Carolina – Chapel Hill, USA) and Cystic Fibrosis Foundation Therapeutics, were generated using purified CFTR as antigen [27]. Recombinant mouse CFTR-NBD1 was produced and purifed as before [67] and injected in two Wistar rats. Animals were injected with 100 µg of antigen with Sigma Adjuvant System at two-week intervals for the first two boosts and with 50 µg of antigen for the third and fourth boosts at a month interval. Hybridoma supernatants were screened by ELISA and Western blot. The resulting monoclonal 3G11 immunoprecipitates CFTR [68] and recognizes aa N396-F405 in the N-terminus of NBD1, was generously provided by Dr. William Balch (Scripps Research Institute, USA) and monoclonal mouse antibodies 2.39.14 and 2.3-5 by Dr. Eric Sorscher (University of Atlanta, Emory, USA). The monoclonal mouse antibody MM13-4 was purchased from Millipore.

Peptides from the first extracellular loop of TMD1 (aa A107-S118), C-terminus of TMD1 (aa S364-K381), ICL4 in TMD2 (aa Q1035-S1049), and the C-terminal part of TMD2 (aa E1172-Q1186) were selected with the AbDesigner algorithm of NIH (https://hpcwebapps.cit.nih.gov/AbDesigner/) and synthesized as described [69]. The purity of synthetic peptides was analyzed by HPLC and mass spectrometry. Synthetic peptides were coupled to rabbit serum albumin (RSA) with the linker Sulfo-*m*-maleimidobenzoyl-*N*-hydroxysuccinimide ester (Sulfo-MBS, Thermo Scientific) in a carrier (1):linker (50):peptide (50) molar ratio. RSA and Sulfo-MBS were dissolved separately in 500 μl of PBS at pH 8.4, mixed and incubated at room temperature for 45 minutes. RSA-sulfo-MBS was desalted on a PD10 column (GE Healthcare) with PBS at pH 7.2, and concentrated to 1 mL (Vivaspin 20, GE Healthcare). Desalted RSA-sulfo-MBS was added to the peptides, which were dissolved in 200 μl of PBS, pH 7.2, and incubated for 2 hours at room temperature. The RSA-coupled peptides were used for antibody production (Pocono rabbit farm & Laboratory, Canadensis, PA). Corrector compound VX-809 and VX-770 (Selleck chemicals) were dissolved in DMSO and stored at −80°C. Proteinase K from Tritirachium album (quality level ELITE) was purchased from Sigma.

### Expression constructs

CFTR single-domain constructs were generated in pBS vectors as previously described [7], and subcloned into pBI-CMV2 using NotI and XhoI. The hCFTR construct in pBI-CMV2 was a kind gift of Linda Millen and Dr. Phil Thomas (University of Texas Southwestern Medical Center, USA). The TMD2-3HA construct was generated by Gibson assembly from pBI-CMV2-CFTR and a gene block consisting of triple HA, with 3xHA positioned at position S898 in ECL4 of CFTR. The TMD1-L-TMD2 construct was also generated by Gibson assembly from pBI-CMV2-1202X, pcDNA3.1-Pgp, and included aa M1-M394 (TMD1) and aa M837-D1202 (TMD2) from CFTR, and aa 626-683 from P-gp in between that acts as a linker. N-terminally truncated constructed were generated from pBI-CMV2-CFTR by PCR. PCR products were then cloned into pBI-CMV2 using AflII and NheI. To retain the same 5’-UTR we also generated a wild-type CFTR with the same cloning strategy. C-terminally truncated constructs were generated from pBI-CMV2-CFTR by PCR and cloned into pBI using NotI and SalI. CFTR mutations were made from pBI-CMV2-CFTR template using high throughput PCR mutagenesis [70]. In short, two fragments are created using the AmpR forward primer in combination with the mutant reverse primer and vice versa and subsequently ligated together using Gibson assembly [71]. A full list of PCR primers is described in the supplementary materials (Table S6).

Construct nomenclature: N-terminal truncation ΔN48 starts with Methionine followed by residues 49 and following. C-terminal truncation K381X has a stop codon instead of K at position 381, and therefore has 380 as the most C-terminal residue.

### Cell culture and transfection

HEK 293T cells were maintained in DMEM supplemented with 10% FBS and 2 mM GlutaMAX (growth medium) and incubated at 37 °C with 5% CO^2^. Cells were seeded onto polylysine-coated 6 cm dishes to reach 70% confluency and transfected using linear 40 kDa polymer polyethylenimine (PEI) as described [24]. After 4 hours, the transfection mix was replaced with growth medium and cells were cultured for 16-20 hours prior to experiments.

### Radioactive pulse and chase

HEK239T cells were used in pulse-chase assays as described [72, 73]. Cells were pulse labeled for 15 minutes with 132 μCi/6 cm dish with EasyTag Express ^35^S Protein Labeling Mix (Perkin Elmer). Radiolabeling was stopped by adding excess, unlabeled 5 μM methionine and 5 μM cysteine. At indicated chase times, cells were washed twice with ice-cold Hanks’ balanced salt solution (HBSS, Life Technologies) and solubilized in ice-cold lysis buffer (20 mM MES, 100 mM NaCl, 30 mM Tris-HCl pH 7.4, 1% Triton X-100) without protease inhibitors. Nuclei were removed by centrifugation for 10 minutes at 4 °C in a microfuge at maximum speed, and the supernatant was subjected to limited proteolysis or immunoprecipitation.

### In-vitro translation and translocation

Target mRNA was prepared by transcribing DNA using T7 RNA polymerase according to the manufacturer’s instructions (Promega). In-vitro translations were done with rabbit reticulocyte lysate (Flexi Rabbit Reticulocyte Lysate System, Promega) and translated CFTR (domains) were translocated and inserted into HEK293-derived microsomes or semi-permeable HEK293T cells as source of ER membrane as described [7]. In brief, target mRNA was added to the reaction mix containing rabbit reticulocyte lysate, 10 μCi/μL EasyTag Express ^35^S Protein Labeling Mix (Perkin Elmer), and membranes. The reaction proceeded at 30°C for 30 minutes, was stopped with 1 mM cycloheximide, and membranes were pelleted through centrifugation at 10,000 x g for 3 minutes at 4°C. Newly translated proteins were retrieved from the pellet fraction by dissolving the membranes in 10 μL KHM (110 mM KOAc, 2 mM Mg(OAc)_2_, 20 mM HEPES pH 7.2) containing 1% Triton X-100 for 10 minutes at 4°C. Samples were either subjected to immunoprecipitation or used directly for SDS-PAGE analysis after adding 2x Laemmli sample buffer.

### Preparation of semi-permeabilized cells

The preparation of semi-permeabilized cells was described before [74]. Confluent HEK293T cells (∼80%) from a 10-cm dish were trypsinized, resuspended in 9 mL ice-cold KHM containing 10 µg/mL Soybean Trypsin Inhibitor (Sigma) and transferred to a 15-mL polypropylene tube. Starting from this step, the cells were kept on ice and centrifugation was done at 4°C. Cells were spun down at 250 x g for 3 min and resuspended in 6 mL ice-cold KHM. To selectively permeabilize the plasma membrane, 40 µg/mL digitonin (Calbiochem) was added and cells were mixed by inversion. After exactly 5 min incubation on ice, permeabilization was stopped by adding 8 mL ice-cold KHM. Cells were immediately spun down, resuspended in 10 mL ice-cold HEPES buffer (50 mM HEPES pH 7.2, 90 mM KOAc), and incubated for 10 min on ice. The semi-permeable cells then were resuspended in 1 mL ice-cold KHM and transferred to a 1.5-mL tube. To check whether permeabilization succeeded, Trypan Blue (Fluka) was added to the cells and the fraction of permeable cells counted by light microscopy. Next, the semi-permeable cells were briefly spun down for 15 seconds at 10,000 x g and resuspended in 100 µL KHM. To degrade endogenous mRNA, 1 mM CaCl_2_ and 10 µg/mL micrococcal nuclease (GE Healthcare) were added and incubated for 12 min at room temperature. To inactivate micrococcal nuclease, 4 mM EGTA was added to chelate calcium. After a final spin down and resuspension in KHM, cells are ready for use in vitro.

### Endoglycosidase H and PNGase F treatment

After in-vitro translation, the membrane fraction containing translocated CFTR was dissolved in 10 µL 100 mM NaAc (pH 5.5) and 1% Triton X-100. 500 U of Endoglycosidase H was added and incubated at 37°C for 1 hour. Samples were used for SDS-PAGE analysis after adding 2x Laemmli sample buffer. PNGase F treatment was performed after immunoprecipitation. The beads were resuspended in PBS containing 0.2% SDS and heated for 5 minutes at 55 °C. Triton was added at a final concentration of 2% to quench the SDS, and 1 U of PNGaseF was added and incubated at 37 °C for 1.5 hours. Samples were used for SDS-PAGE analysis after adding 5X sample buffer.

### Limited proteolysis

Limited proteolysis was performed as described [7, 23]. In brief, lysates were treated with 25 μg/mL Proteinase K (Sigma-Aldrich) for 15 minutes on ice. Proteolysis was stopped by mixing equal volumes of lysis buffer supplemented with 2 mM PMSF and 2 μg/mL CLAP (chymostatin, leupeptin, antipain and pepstatin (Sigma-Aldrich)) with the lysates. Protein aggregates were pelleted by 16,000 x g centrifugation for 5 minutes at 4°C, and the supernatant was used for immunoprecipitation.

### Immunoprecipitation

Antibodies against CFTR were pre-incubated with protein-A or protein-G Sepharose beads (GE Healthcare) for 15 minutes at 4°C before adding protease-treated or non-treated lysates. The lysates were added to the antibody-beads mixtures and incubated at 4°C for either 3 hours or overnight. The beads were washed twice for 15 minutes at room temperature. A list of immunoprecipitation conditions is provided in the supplementary materials (Table S7). Beads were resuspended in 10 μl 10 mM Tris-HCl pH 6.8 containing 1 mM EDTA, and immune complexes were eluted by adding 10 μl 2x reducing Laemmli sample buffer (final concentration: 200 mM Tris-HCl pH 6.8, 3% SDS, 10% glycerol, 1 mM EDTA, 0.004% bromophenol blue, and 25 mM DTT) and heating for 5 min at 55 °C.

### SDS-PAGE and autoradiography

Samples generated by in-vitro translation or fragments generated by limited proteolysis were resolved by 12% SDS-PAGE, whereas all other samples were resolved by 7.5 or 10% SDS-PAGE, as indicated in the legends. Gels were dried and exposed to super-resolution phosphor screens (Fuji Film) for quantifications, or to Kodak MR films for manuscript figure images. Signals from screens were visualized with a Typhoon FLA-7000 scanner (GE Healthcare Life Science) and quantified with ImageQuantTL software (GE Healthcare Life Science).

### Structural analysis

The helical propensity of domain sequences was determined using secondary structure prediction algorithms via JPred v4 [75]. Information of secondary structures of NBD1 and NBD2 of CFTR were taken from PDB (5UAK). Images of protein structures were created using UCSF Chimera [76].

## ACKNOWLEDGEMENTS

We thank Dr. William Balch (Scripps Research, La Jolla CA USA) for kindly providing the 3G11 antibody, Dr. Angus Nairn (Yale, New Haven CT USA) and Dr. Hugo de Jonge (Erasmus Medical Center, Rotterdam NL) for the G449 antibody, Dr. John Riordan (University of North Carolina, Chapel Hill NC USA), and the Cystic Fibrosis Foundation (CFF) for antibodies 570, 217 and 596, and Dr. Eric Sorscher (Emory University, Atlanta GA USA) for the 2.3-5 and 2.39.14 antibodies. We thank Linda Millen and Dr. Phil Thomas (University of Texas Southwestern Medical Center, Dallas TX USA) for kindly providing the hCFTR construct in pBI-CMV2. We thank Laurens Kooijman MSc for designing the peptides, and Dr. Maarten Egmond and Ing. Ruud Cox for supervising peptide synthesis. We are grateful to Dr. Joseline Houwman for discussions, suggestions, critical reading, the artwork, and editing and thank all members of the Braakman-Van der Sluijs lab for fruitful discussions. This work was funded by Cystic Fibrosis Foundation, Netherlands Cystic Fibrosis Foundation, Stichting Zeldzame Ziekte Fonds via Stichting Muco & Friends and Netherlands Organization for Health Research and Development-Zon-MW Top (to IB), Dutch Research Council (to MM), European Union Horizon 2020 and long term EMBO postdoctoral fellowship (to PS).

## STATEMENTS & DECLARATIONS

### Author Contributions

Conceptualization: Jisu Im, Marjolein Mijnders, Ineke Braakman; Formal analysis and investigation: Jisu Im, Tamara Hillenaar, Marcel van Willigen, Marjolein Mijnders, Hui Ying Yeoh; Writing - original draft preparation: Jisu Im; Writing - review and editing: Jisu Im, Peter van der Sluijs, Ineke Braakman; Funding acquisition: Ineke Braakman, Peter van der Sluijs, Marjolein Mijnders; Resources: Hui Ying Yeoh, Priyanka Sahasrabudhe; Supervision: Peter van der Sluijs, Ineke Braakman

### Data Availability

The datasets generated during and/or analysed during the current study are not publicly available due to the complexity of the data but are available from the corresponding author on reasonable request.

