## Supplementary material for "ABC-transporter CFTR folds with high fidelity through a modular, stepwise pathway": Suppl. Method and Tables

### SUPPLEMENTARY TEXT

#### Workflow for identification of domain-specific protease-resistant fragments

To identify the regions in CFTR that are protected or de-protected from the protease during folding we determined the identity of each proteolytic fragment, i.e. the cleavage sites used to generate the domain-specific protease-resistant fragments. We mapped the presence of antigenic epitopes and glycans in the fragments, electrophoretic mobility shifts in the fragments in missense-mutated CFTR and truncated CFTR constructs, and considered helical propensity and consensus cleavage sites of the protease to locate the cleavage sites even further. With this information we identified each cleavage site, down to just a few amino acids for most fragments, a treasure trove for many future studies.

We set out to determine the identity of each proteolytic fragment and started by mapping the presence of antigenic epitopes (Table S1, Figures 3a, 5a, 7c) and glycans in each fragment. In a next step we truncated CFTR domains or parts and analyzed resulting electrophoretic mobility shifts in SDS-PAGE, which allowed determination of several fragment boundaries already (Tables 1, S3-5 and Figures 3, 5e, 7d). Last experimental dataset was the mapping of fragment mobility shifts (or lack thereof) in point mutants and in truncated CFTR (Tables 1, S3-5 and Figures 3d, 5, 8b). These results, combined with literature on proteinase K consensus cleavage sites (Table S2) and alpha-helical-propensity predictions, allowed determination of the amino-acid sequences (up to a single likely residue) that constituted the N-terminal and C-terminal cleavage sites for each fragment (Tables 1, S3-5 and Figures 4a, 6a and 7a-b).

We found that the electrophoretic mobility of fragments (especially from TMD) was not dictated by size/mass alone, but strongly by charge and hydrophobicity as well. For instance, the electrophoretic mobility of fragment T1a suggested a size of ~19 kDa, but this was too small to fit within the experimentally determined fragment boundaries (minimal 23 kDa). We therefore did not use this as a strict parameter to determine cleavage sites.

Our analysis depends fully on the specificity of proteinase K. Other proteases such as trypsin, chymotrypsin, endoGlu-C, and thermolysin are useful to confirm our findings, but the strength of proteinase K is its low specificity, with possible cleavage sites all throughout the CFTR sequence. This implies that protection from cleavage can be caused only by conformational or external factors. Proteases select against helical sequences [1, 2], which we used by predicting helical propensity of cleavage stretches using Jpred (http://www.compbio.dundee.ac.uk/jpred/). Cleavage sites are shielded by interaction with polypeptide chains, either within the molecule (domain folding and domain assembly), or from another molecule, for instance by assembly with other subunits in an oligomer or by a (transient) interaction with another protein such as a chaperone. Our analysis in Triton X-100 favors intramolecular interactions and tightly associating oligomers and disfavors chaperone associations. We therefore conclude that proteolytic changes are caused by CFTR folding, although we cannot exclude intermolecular interactions.

The consensus cleavage sites of proteinase K have been well established but unfortunately some ambiguity remains. Proteinase K preferentially cleaves at the carboxyl side of residues A, Q, M, C, W, S, F or L (Table S2) [3-7] located in accessible loops and not in secondary structure, especially α- 221 helices [1, 2]. Cleavage of residues that are followed by small amino acids such as alanine or glycine is slightly preferred whereas cleavage is inhibited when the residue is followed by a proline and slightly reduced when followed by valine [8]. Also, cleavage is more efficient when the cleavage site is flanked by at least 1 to 3 residues N- or C-terminal [4, 8].

The antigenic epitopes and truncated constructs provided the primary information on the preferred cleavage sites and are written in Tables 1, S3-5 (and Figures 4a, 6a and 7a-b). Next are the consensus proteinase-K sites, and the electrophoretic mobility shifts caused by missense mutations. The reasoning for the N- and C-terminal boundaries of each fragment is in Tables S3-5 and shown in Figures 4a, 6a and 7a-b.

A missense mutation can influence proteolysis by three mechanisms:

1. Removal or addition of a consensus proteinase-K cleavage site,

2. Change of electrophoretic mobility by charge or changed SDS binding, and
3. Change of conformation or interactions with other proteins, leading to different shielding of consensus sites.

Ad 1. Change of a cleavage site may go unnoticed when a cluster of sites is being used by the protease, implying that we cannot conclude that a lack of change equates a lack of cleavage of the target or mutated residue. Exceptions to this are the few instances where a single residue appears to be the protease target rather than an amino-acid cluster. A change in proteolytic fragment caused by a mutated cleavage site does provide positive information.

Ad 2. Electrophoretic mobility of proteins in SDS-PAGE [9] in principle is determined by their mass because of their inherent capacity to bind 1.4 g of SDS per g polypeptide, independent of amino acid composition. As SDS adds 2 negative charges it in principle overrules any charge the protein may have. These ideal conditions often are not met, for instance in hydrophobic proteins, where SDS binding may be higher or lower, or in negatively charged proteins such as all calcium-binding chaperones in the endoplasmic reticulum, where SDS binding is lower, leading to decreased mobility and overestimation of their masses. Analyzing smaller constructs, the individual CFTR domains and proteolytic fragments, we noticed that a charge change in the fragment did contribute to a mobility change, either up or down, by a combination of its own charge and the changed SDS-binding. On top of the charge changes we have noticed electrophoretic mobility changes unrelated to charge, such as in I539T [10]. In summary, this parameter, while important to notice and include in the analysis, does not provide rigorous information on proteolytic fragments. Mobility appeared most useful to establish whether the mutated residues were present in a fragment.

Ad 3. Limited proteolysis classically has been used for this parameter: to determine folded state or domain boundaries of proteins [1]. In Cleveland analysis [11] it is used as unique protein identifier, on the premise that each native protein has its own sequence and hence unique proteolytic pattern. We have used the approach and found a surprising capacity to zoom into the used cleavage sites. This is fortunate as mass spectrometry so far has had limited success in determining identities of these subpicomole quantities of radiolabeled fragments. Equating proteolysis to folded state of CFTR is only possible when other proteins are not interacting and shielding protease sites. As Triton X-100 dissociates most chaperone-client and many other complexes, we limited our conclusions here to intramolecular shielding, i.e. folding.

Following this list of parameters for the N-terminus and C-terminus of each proteolytic fragment, described in Tables S3-5 and visualized in Figures 4a, 6a and 7a-b, we identified each fragment with rather precise boundaries (Table 1).

### SUPPLEMENTARY FIGURE LEGENDS

**Figure S1. Glycosylation of TMD2 in vitro and CFTR in cells, related to Figure 1.**
**(a)** TMD2 was translated in vitro in the presence of HEK293 microsomes. Membranes were pelleted, digested with Endo H where indicated, and resolved by 12% SDS-PAGE. TMD2gg: core-glycosylated TMD2. **(b)** HEK293T cells expressing CFTR were pulse-labeled for 15 minutes and lysed immediately or after a chase of 2 hours, immunoprecipitated using MrPink antibody and resolved by 10% SDS-PAGE.

**Figure S2. PNGase F deglycosylates core and complex glycosylated CFTR, related to Figure 5**
HEK293T cells expressing full-length CFTR were pulse-labeled for 15 minutes and lysed immediately or after a chase of 2 hours. CFTR was immunoprecipitated with MrPink, subjected to PNGase F treatment, and resolved by 7.5% SDS-PAGE.

**Figure S3. Protease resistant fragments shift downwards in N-terminally truncated CFTR, related to Figure 3d-g.**
**(a)** HEK293T cells expressing N-terminal CFTR truncations ΔN48 to ΔN76 were pulse-labeled for 15 minutes and lysed immediately. CFTR variants were immunoprecipitated with MrPink and analyzed by 7.5% SDS-PAGE. Digests are shown in Figure **3b**. **(b)** Same as **(a)** but now with N-terminal CFTR truncations ΔN47 to ΔN51. Digests are shown in Figure **3c**. **(c)** same as **(a)** but with C-terminal truncations K381X to E395X. Digests are shown in Figure **3d**. **(d)** same as **(a)** but for N-terminal truncations ΔN35 to ΔN45, and after a chase of 1 hour. Digests are shown in Figure **3f**. **(e)** same as **(d)** but now for N-terminal truncations ΔN2 to ΔN5. Digests are shown in Figure **3g**. Experiments in panels **(d)** and **(e)** were done in the presence of VX-770 (3 μM) and/or VX-809 (3 μM) as indicated.

**Figure S4. Protease resistant fragments shift in CFTR mutants, related to Figure 5b-e.**
**(a)** HEK293T cells expressing CFTR mutants N901K-V905K were pulse labeled for 15 minutes and lysed after a 2-hour chase. Detergent lysates were immunoprecipitated with MrPink or subjected to limited proteolysis with 25 μg/mL proteinase K and immunoprecipitated with TMD2C. Top panel represents non-digested material resolved by 7.5% SDS-PAGE and the bottom panel shows the

fragments generated by limited proteolysis and resolved by 12% SDS-PAGE. **(b)** same as **(a)** but with mutants L957K to G970K and 0-hour chase. Dotted red line through T2b fragment is for visual aid. **(c)** HEK293T cells expressing CFTR mutants S1058K to L1065K were pulse labeled for 15 minutes and lysed immediately. Detergent lysates were immunoprecipitated with MrPink and resolved by 7.5% SDS-PAGE. **(d)** same as **(c)** but with mutants M961K to L967K. **(e)** same as **(c)** but with mutants T908K to S912K and after a chase of 2h. **(f)** same as **(bc)** with C-terminal truncations N1184X to M1191X. **(g)** same as **(b)** but with mutants V1190X to S1196X. Dotted red line through T2b fragment is for visual aid. Digests of panels **c-f** are shown in Figure **5b-e**.

### SUPPLEMENTARY TABLES

#### Table S1. CFTR Antigenic epitopes

*Antibodies generated against their epitope peptide.

| **Antibody** | **Domain** | **Epitope** (residues) |
|---|---|---|
| MM13-4 | TMD1, N-terminus | G27-L34 [12] |
| E1-22 | TMD1, ECL1 | A107-S118* [13] |
| TMD1C | TMD1, C-terminus | S364-K381* |
| 3G11 | NBD1, N-subdomain | N396-F405 (personal communication W. Balch) [13] |
| MrPink | NBD1, whole | Multiple in NBD1 [10] |
| G449/G450 | R region & NBD1 C-subdomain | F653-K716* [14] |
| 570 | R region | P731-S742 [12] |
| 217 | R region | I807-E819 [12] |
| I4N | TMD2, ICL4 | Q1035-S1049* |
| TMD2C | TMD2, C-terminus | E1172-Q1186* |
| 596 | NBD2, N-subdomain | W1204-T1211 [12] |
| 2-39.14 | NBD2, C-subdomain | E1371-R1385 (E. Sorscher) |
| 2-3.5 | NBD2, C-subdomain | V1379-T1387 (E. Sorscher) |

**Table S2. Proteinase K consensus cleavage studies**

Summary of literature describing which amino acids (displayed as one-letter code) Proteinase K preferentially cleaves. Amino acids that were not or barely cleaved are in regular font. This was generally consistent throughout all studies. We chose to use the residues in bold as preferentially cleaved by proteinase K, mostly based on the study performed by Whittaker et al., which is the most rigorous and complete one. These are colored blue in Figures 4 and 6. Results on tyrosine (Y) were too variable to consider Y a consensus cleavage site.

| Study | Preferential cleavage of amino acids |
|---|---|
| [7] | **A>Q>M>C>W>S>F>L**>H>N>E=V>T>Y> K=G>R>D>I>P |
| [4] | **W**>Y>**F>L>A**> K>R>V>G and K>F>A |
| [3] | Y>**A=L**> R |
| [5] | Y>**F>L**> P=K=R=G |
| [6] | **Q-C-F-L**-Y-N-H-S-E-G> V=A=R=T=P=K |

**Table S3. Cleavage site identification of TMD1**

Indirect, supporting results are in *italics*, conclusion is in **bold**

| Fragment | N- or C-terminus | Results | Reference |
|---|---|---|---|
| T1a | N | • Does not contain MM13-4 epitope (aa 27-34). | Fig. 3a |
| | | • Shifts on SDS-PA gel in N-terminal truncations: ΔN48 (no shift), ΔN55 (shift down) | Fig. 3b |
| | | • Shifts on SDS-PA gel in N-terminal truncations: ΔN47 (no shift), ΔN48 (no shift), ΔN49 (no shift), ΔN50 (shift down), ΔN51 (shift down). | Fig. 5b in [15] |
| | | • *Lh2 (aa46-63) has relatively low helical propensity.* | [16] |
| | | • *Proteinase-K cleavage sites between aa46-55 are A46, L49, S50 and L53.* | Table S2 |
| | | • **N-terminal cleavage site is L49** | |
| | C | • Contains ECL1 epitope (aa107-118), does not contain TMD1C epitope (aa362-381) | Fig. 3a |
| | | • Shifts on SDS-PA gel in C-terminal truncations: 249X (shift down), 307X (no shift) | Fig. 4b in [15] |
| | | • No shift in C-terminal truncations (357X, 366X, 376X, 386X, 395X) | Fig. 7b in [15] |
| | | • Shifts on SDS-PA gel in C-terminal truncations: 254X (shift down), 255X (shift down), 256X (no shift), 257X (no shift), 258X (shift up), 259X (no shift), 260X (no shift), 261X (no shift) | Fig. 5c in [15] |
| | | • *ICL2 has a high helical propensity, apart from aa275-278: which is the loop after the ICL2 coupling helix.* | [16] |
| | | • **C-terminal cleavage site is S256** | |
| T1c | N | • Does not contain MM13-4 epitope (aa27-34). | Fig. 3a |
| | | • Shifts on SDS-PA gel in ΔN55 (shift down). | Fig. 3b |
| | | • Shifts in N-terminus truncations: ΔN47 (no shift), ΔN48 (no shift), ΔN49 (no shift), ΔN50 (down), ΔN51 (down). | Fig. 3c |
| | | • *Lh2 (aa46-63) relatively low helical propensity.* | [16] |
| | | • *A46, L49, S50 and L53 are only Proteinase-K consensus cleavage sites between aa46-55.* | Table S2 |
| | | • **N-terminal cleavage site is L49** | |
| | C | • Contains TMD1C epitope (aa362-381) does not contain 3G11 epitope (aa396-405). | Fig. 3a |
| | | • Shift changes in C-terminal truncations: 381X (shift down), 382X (shift down), 383X (shift down), 384X (no shift), 385X (no shift) and 395X (no shift) | Fig. 3f |
| | | • *aa 381-396 forms a flexible loop.* | [17] |
| | | • *Proteinase-K consensus sites between aa381-396 are the following: L383, L387 and M394. M394 is located in a β-sheet so less likely to be cleaved.* | Table S2 |
| | | • **C-terminal cleavage site is L383** | |
| T1d | N | • No MM13-4 epitope (aa 27-34). | Fig. 3a |
| | | • Shift changes in N-terminal truncations: ΔN35 (no shift), ΔN45 (shift down). | Fig. 3e |
| | | • Shifts in N-terminal truncations: ΔN35 (no shift), ΔN37 (shift down), ΔN40 (shift down), ΔN42 (shift down), ΔN44 (shift down) ΔN45 (shift down). | Fig. 3f |

| Fragment | N- or C-terminus | Results | Reference |
|---|---|---|---|
| | | • *aa36-45 form a flexible linker*<br>• *S36, Q39 and S45 are only Proteinase-K consensus sites between aa30-45 (S42 is followed by V43 and therefore less likely cleaved).*<br>• **N-terminal cleavage site is S35** | [17]<br>Table S2 |
| | C | • Contains TMD1C epitope (aa362-381) does not contain 3G11 epitope (aa396-405).<br>• *aa 381-396 forms a flexible loop.*<br>• *Proteinase-K consensus sites between aa381-396 are the following: L383, L387 and M394. M394 is located in a β-sheet so less likely to be cleaved. L383 is the cleaved residue in the more open, unfolded, protease-sensitive CFTR yielding T1c.*<br>• **C-terminal cleavage site is most likely L383** | Fig. 3a<br><br>[17]<br>Table S2 |
| T1e | N | • Contains MM13-4 epitope (aa 27-34)<br>• Shift changes in N-terminal truncations: ΔN16 (present), ΔN18 (present), ΔN20 (vague), ΔN35 (down)<br>• *Proteinase-K consensus sites between aa17-27 are F17, S18, W19 and L24. F17 most likely cleaved.*<br>• **N-terminal cleavage site is most likely F16** | Fig. 3a<br>Fig. 3e<br><br>Table S2 |
| | C | • Identical to T1d.<br>• **C-terminal cleavage site is most likely L383** | |
| T1f | N | • Contains MM13-4 epitope (aa27-34).<br>• Shift changes in N-terminal truncations: ΔN4 (down)<br>• Shifts in N-terminal truncations: ΔN2 (down), ΔN3 (down), ΔN4 (down), ΔN5 (down).<br>• **No N-terminal cleavage site, starts with M1** | Fig. 3a<br>Fig. 3e<br>Fig. 3g |
| | C | • Identical to T1d.<br>• **C-terminal cleavage site is most likely L383** | |

**Table S4. Cleavage site identification of TMD2**

Indirect results are in *italics*, conclusion is in **bold**

| Fragment | N- or C-terminus | Results | Reference |
|---|---|---|---|
| T2a | N | • Glycans not present.<br>• Contains TMD2C epitope (aa1172-1186), does not contain ICL4 epitope (aa1035-1049).<br>• *Shifts in ICL4 mutations on SDS-PA gel: S1058K (no shift), L1059K (absent, smaller doublet), K1060R (no shift), G1061K (shift up), L1062K (shift up, smaller doublet), W1063K (shift up, smaller doublet), T1064K (shift up) and L1065K (no shift).*<br>• *Helical propensity is low on both sides of the ICL4 coupling helix.*<br>• *Proteinase-K consensus sites in ICL4 region are the following: L1059, L1062, W1063, L1065, A1067, F1068. All residues here are relatively high on preferential cleavage of Proteinase-K.*<br>• **N-terminal cleavage site is L1059** | Fig. 5a<br>Fig. 2e,f, 5a<br>Fig. 5b<br>[16]<br>Table S2 |
| | C | • Contains TMD2C epitope (aa1172-1186), does not contain 596 epitope (aa1204-1211)<br>• Shifts in C-terminal truncations on SDS-PA gel: M1191X (no shift), V1190X (not detectable), K1189X (shift down), S1188X (shift down), L1187X (shift down), Q1186X (shift down), G1185X (shift down) and N1184X (shift down)<br>• *Proteinase-K consensus sites between aa1186-1204 are L1187, S1188, M1191 and W1204. W1204 is likely blocked by P1205.*<br>• **C-terminal cleavage site is M1191** | Fig. 5a<br>Fig. 5e<br>Table S2 |
| T2b | N | • Glycans not present.<br>• Contains TMD2C epitope (aa1172-1186) and contains ICL4 epitope (aa1035-1049)<br>• *Helical propensity higher in N-terminal than C-terminal of ICL3 coupling helix.*<br>• *Shift changes in ICL3 mutants on SDS-PA gel shown in C-terminal half of ICL3: M961K (no shift), S962K (no shift), T963K (no shift), L964K (shift up), N965K (no shift), T966K (doublet), L967K (doublet), A969K (doublet).*<br>• *Proteinase-K consensus sites in C-terminal region of ICL3 are the following: L964, L967, A969.*<br>• **N-terminal cleavage site is L964** | Fig. 5a<br>Fig. 2e, 2f, 5a<br>[16]<br>Fig. 5c, S4b<br>Table S2 |
| | C | • Contains TMD2C epitope (aa1172-1186), does not contain 596 epitope (aa1204-1211)<br>• Shift changes in C-terminal truncations: S1196X (no shift), N1195X (no shift), E1194X (no shift), I1193X (no shift), I1192X (no shift), M1191X (no shift), V1190X (no shift),K1189X (shift down), S1188X (shift down), L1187X (shift down), Q1186X (shift down), G1185X (shift down) and N1184X (shift down)<br>• *Proteinase-K consensus sites between aa1186-1204 are L1187, S1188, M1191 and W1204. W1204 is likely blocked by P1205.*<br>• **C-terminal cleavage site is M1191** | Fig. 5a<br>Fig. 5e, S4g<br>Table S2 |

| Fragment | N- or C-terminus | Results | Reference |
|---|---|---|---|
| T2c | N | • Glycans not present.<br>• Contains TMD2C epitope (aa1172-1186) and contains ICL4 epitope (aa1035-1049).<br>• T2c > T2b<br>• *Shift changes in ECL4 charge mutants: N901K (no shift), S902K (no shift), Y903K (no shift), V905K (no shift), T910K (no shift), S911K (shift up), S912K (shift up).*<br>• *Proteinase-K consensus sites in ECL4 region are the following: S902, A904, S909, S911, S912.*<br>• **N-terminal cleavage site is S909** | Fig. 5a<br>Fig. 2e, 2f, 5a<br><br>Fig. 5d, S4a<br><br>Table S2 |
| | C | • Identical to T2a and T2b<br>• **C-terminal cleavage site is most likely M1191** | |

**Table S5. Cleavage site identification of NBD2**

Indirect results are in *italics*, conclusion is in **bold**

| Fragment | N- or C-terminus | Results | Reference |
|---|---|---|---|
| N2a | N | • Contains 596 epitope (aa1204-1211), does not contain TMD2C epitope (aa1172-1186) | Fig. 7c |
| | | • *M1191 is cleavage site for TMD2 fragments. Only other Proteinase-K consensus sites between aa1186-1204 are L1187, S1188, M1191 and W1204. W1204 is likely blocked by P1205. L1187, S1188 are options.* | Table S2 |
| | | • **N-terminal cleavage site is M1191** | |
| | C | • Contains 2-39.14 epitope (aa1370-1385) and 2-3.5 epitope (aa1379-1387) | Fig. 7c |
| | | • Shift changes in C-terminal truncations: F1437X (shift down), R1438X (shift down), Q1439X (shift down), A1440X (shift down), I1141X (no shift), S1442X (no shift). | Fig. 7d |
| | | • *Size of the fragment and similarity to NBD1 sµggests N2a includes almost of all NBD2 lacking the flexible C-terminus (aa1436-C term).* | |
| | | • **C-terminal cleavage site is Q1439** | |

**Table S6. Primers used for generation of constructs used in this study**

| Construct Name | Primer Sequence (5' to 3') |
|---|---|
| CFTR-TMD1 (CFTR 395X) | F: GGAATGCAGATGAGAATAGC<br>R: CCCTCGAGTCTACATCACTACTTCTGTAGTCG |
| CFTR-TMD2 (837-1202) | F: GCTCTAGAATGGAGAGCATACCAGCAGTGAC<br>R: CCCTCGAGCTAATCTTTCTTCACGTGTGAATTCTC |
| TMD2-3HA | F: AGAAATAACAGCTATGCAGTG<br>R: ACTATGAGTACTATTCCC |
| | Gene block:<br>GGGAATAGTACTCATAGTGGTACCTACCCTTACGATGTTCCTGACTAT<br>GCGGGCTATCCGTACGACGTCCCGGACTATGCAGGATCCTATCCATAT<br>GATGTGCCAGATTACGCTACTAGTAGAAATAACAGCTATGCAGTG |
| TMD1-L-TMD2 | F: AGAATTCACACGTGAAGAAAGATTGACGGCCGCATCGATAAG<br>R: CCTGCTGTCTGCATTGTGACCATCACTACTTCTGTAGTCGTTAAGTT |
| | F: CGACTACAGAAGTAGTGATGGTCACAATGCAGACAGCAGG<br>R: AGGTATGTGTTCCATGTAGTAAGCTTTCTGTCTTGGGCTT |
| | F: GCTTATCGATGCGGCCGTCAATCTTTCTTCACGTGTGAATTCT<br>R: AAGCCCAAGACAGAAAGCTTACTACATGGAACACATACCTT |
| Wild-type (- 5'-UTR) | F: ATAGCTAGCACCATGCAGAGGTCGCCTCTGGAAAAG<br>R: GGCTGAGCTATTGAAGTATCTCACATAGGCTG |
| ΔN2 | F: ATAGCTAGCACCATGAGGTCGCCTCTGGAAAAGG<br>R: CCGCTCGAGCTAATATTCCAATGTCTTATA |
| ΔN3 | F: ATAGCTAGCACCATGTCGCCTCTGGAAAAGGCC<br>R: CCGCTCGAGCTAATATTCCAATGTCTTATA |
| ΔN4 | F: ATAGCTAGCACCATGCCTCTGGAAAAGGCCAGCGTTGTC<br>R: GGCTGAGCTATTGAAGTATCTCACATAGGCTG |
| ΔN5 | F: ATAGCTAGCACCATGCTGGAAAAGGCCAGCGTTGTC<br>R: GGCTGAGCTATTGAAGTATCTCACATAGGCTG |
| ΔN8 | F: ATAGCTAGCACCATGGCCAGCGTTGTCTCCAAAC<br>R: GGCTGAGCTATTGAAGTATCTCACATAGGCTG |
| ΔN10 | F: ATAGCTAGCACCATGGTTGTCTCCAAACTTTTTTTCAGCTG<br>R: GGCTGAGCTATTGAAGTATCTCACATAGGCTG |
| ΔN12 | F: ATAGCTAGCACCATGTCCAAACTTTTTTTCAGCTGGACCAG<br>R: GGCTGAGCTATTGAAGTATCTCACATAGGCTG |
| ΔN16 | F: ATAGCTAGCACCATGTTCAGCTGGACCAGACCAATTTTG<br>R: GGCTGAGCTATTGAAGTATCTCACATAGGCTG |
| ΔN18 | F: ATAGCTAGCACCATGTGGACCAGACCAATTTTGAGG<br>R: GGCTGAGCTATTGAAGTATCTCACATAGGCTG |
| ΔN20 | F: ATAGCTAGCACCATGAGACCAATTTTGAGGAAAGGATAC<br>R: GGCTGAGCTATTGAAGTATCTCACATAGGCTG |
| ΔN35 | F: ATAGCTAGCACCATGGACATATACCAAATCCCTTCTGTT<br>R: GGCTGAGCTATTGAAGTATCTCACATAGGCTG |
| ΔN37 | F: ATAGCTAGCACCATGTACCAAATCCCTTCTGTTGATTCTG<br>R: CCGCTCGAGCTAATATTCCAATGTCTTATA |
| ΔN40 | F: ATAGCTAGCACCATGCCTTCTGTTGATTCTGCTGACA<br>R: CCGCTCGAGCTAATATTCCAATGTCTTATA |
| ΔN42 | F: ATAGCTAGCACCATGGTTGATTCTGCTGACAATCTATCTGAA<br>R: CCGCTCGAGCTAATATTCCAATGTCTTATA |
| ΔN45 | F: ATAGCTAGCACCATGGCTGACAATCTATCTGAAAAATTGG<br>R: GGCTGAGCTATTGAAGTATCTCACATAGGCTG |
| ΔN47 | F: ATAGCTAGCACCATGAATCTATCTGAAAAATTGGAAAGAGAATGGG<br>R: GGCTGAGCTATTGAAGTATCTCACATAGGCTG |
| ΔN48 | F: ATAGCTAGCACCATGCTATCTGAAAAATTGGAAAGAGAATG<br>R: GGCTGAGCTATTGAAGTATCTCACATAGGCTG |
| ΔN49 | F: ATAGCTAGCACCATGTCTGAAAAATTGGAAAGAGAATGGGAT<br>R: GGCTGAGCTATTGAAGTATCTCACATAGGCTG |
| ΔN50 | F: ATAGCTAGCACCATGGAAAAATTGGAAAGAGAATGGGATAGAGAG<br>R: GGCTGAGCTATTGAAGTATCTCACATAGGCTG |

| | |
|---|---|
| ΔN51 | F: ATAGCTAGCACCATGAAATTGGAAAGAGAATGGGATAGAGAG<br>R: GGCTGAGCTATTGAAGTATCTCACATAGGCTG |
| ΔN55 | F: ATAGCTAGCACCATGGAATGGGATAGAGAGCTGGCTTC<br>R: GGCTGAGCTATTGAAGTATCTCACATAGGCTG |
| ΔN65 | F: ATAGCTAGCACCATGAATCCTAAACTCATTAATGCCCTT<br>R: GGCTGAGCTATTGAAGTATCTCACATAGGCTG |
| ΔN68 | F: ATAGCTAGCACCATGCTCATTAATGCCCTTCGGCG<br>R: GGCTGAGCTATTGAAGTATCTCACATAGGCTG |
| ΔN76 | F: ATAGCTAGCACCATGTTTTTCTGGAGATTTATGTTCTATGG<br>R: GGCTGAGCTATTGAAGTATCTCACATAGGCTG |
| 381X | F: CGCAAATGGGCGGTAGGCGTG<br>R: CCGTCGACTTAATATTCTTGCTTTTGTAAGAAATCCTGTATTTTG |
| 382X | F: CGCAAATGGGCGGTAGGCGTG<br>R: CCGTCGACTTACTTATATTCTTGCTTTTGTAAGAAATCCTGTA |
| 383X | F: CGCAAATGGGCGGTAGGCGTG<br>R: CCGTCGACTTATGTCTTATATTCTTGCTTTTGTAAGAAATCCTG |
| 384X | F: CGCAAATGGGCGGTAGGCGTG<br>R: CCGTCGACTTACAATGTCTTATATTCTTGCTTTTGTAAGAAATCC |
| 385X | F: CGCAAATGGGCGGTAGGCGTG<br>R: CCGTCGACTTATTATTCCAATGTCTTATATTCTTGCTTTTGTAAGAAA |
| 395X | F: CGCAAATGGGCGGTAGGCGTG<br>R: CCGTCGACTTACATCACTACTTCTGTAGTCGTTAAGTTATATTC |
| N1184X | F: CGCAAATGGGCGGTAGGCGTG<br>R: GGCGTCGACCTACTTGTATGGTTTGGTTGACTTGG |
| G1185X | F: CGCAAATGGGCGGTAGGCGTG<br>R: GGCGTCGACCTAATTCTTGTATGGTTTGGTTGACTTG |
| Q1186X | F: CGCAAATGGGCGGTAGGCGTG<br>R: GGCGTCGACCTAGCCATTCTTGTATGGTTTGGTTG |
| L1187X | F: CGCAAATGGGCGGTAGGCGTG<br>R: GGCGTCGACCTATTGGCCATTCTTGTATGGTTTGG |
| S1188X | F: CGCAAATGGGCGGTAGGCGTG<br>R: GGCGTCGACCTAGAGTTGGCCATTCTTGTATGG |
| K1189X | F: CGCAAATGGGCGGTAGGCGTG<br>R: GGCGTCGACCTACGAGAGTTGGCCATTCTTG |
| V1190X | F: CGCAAATGGGCGGTAGGCGTG<br>R: GGCGTCGACCTATTTCGAGAGTTGGCCATTCTTG |
| M1191X | F: CGCAAATGGGCGGTAGGCGTG<br>R: GGCGTCGACCTAAACTTTCGAGAGTTGGCC |
| I1192X | F: CGCAAATGGGCGGTAGGCGTG<br>R: GGCGTCGACCTACATAACTTTCGAGAGTTGGCC |
| I1193X | F: CGCAAATGGGCGGTAGGCGTG<br>R: GGCGTCGACCTAAATCATAACTTTCGAGAGTTGGC |
| E1194X | F: CGCAAATGGGCGGTAGGCGTG<br>R: GGCGTCGACCTAAATAATCATAACTTTCGAGAGTTGG |
| N1195X | F: CGCAAATGGGCGGTAGGCGTG<br>R: GGCGTCGACCTACTCAATAATCATAACTTTCGAGAGTTG |
| S1196X | F: CGCAAATGGGCGGTAGGCGTG<br>R: GGCGTCGACCTAATTCTCAATAATCATAACTTTCGAGAG |
| F1437X | F: CGCAAATGGGCGGTAGGCGTG<br>R: GGCGTCGACCTAGAGGCTCCTCTCGTTCAGCAG |
| R1438X | F: CGCAAATGGGCGGTAGGCGTG<br>R: GGCGTCGACCTAGAAGAGGCTCCTCTCGTTCAG |
| Q1439X | F: CGCAAATGGGCGGTAGGCGTG<br>R: GGCGTCGACCTACCGGAAGAGGCTCCTCTCGTTC |
| A1440X | F: CGCAAATGGGCGGTAGGCGTG<br>R: GGCGTCGACCTATTGCCGGAAGAGGCTCCTCTC |
| I1441X | F: CGCAAATGGGCGGTAGGCGTG<br>R: GGCGTCGACCTAGGCTTGCCGGAAGAGGCTCC |
| S1442X | F: CGCAAATGGGCGGTAGGCGTG |

| | |
|---|---|
| | R: GGCGTCGACCTAGATGGCTTGCCGGAAGAGGC |
| AmpR | F: AGCATTTATCAGGGTTATTGTCTCATGAGC<br>R: TAACCCTGATAAATGCTTCAATAATATTGAAAAAGGAAGAGT |
| L957K | F: AAATGTTACATTCTGTTAAGCAAGCACCTATGTCAAC<br>R:GTTGACATAGGTGCTTGCTTAACAGAATGTAACATTT |
| Q958K | F: GTTACATTCTGTTCTTAAAGCACCTATGTCAAC<br>R: GTTGACATAGGTGCTTTAAGAACAGAATGTAAC |
| P960K | F: CTGTTCTTCAAGCAAAGATGTCAACCCTCAAC<br>R: GTTGAGGGTTGACATCTTTGCTTGAAGAACAG |
| M961K | F: CTTCAAGCACCTAAGTCAACCCTCAAC<br>R: GTTGAGGGTTGACTTAGGTGCTTGAAG |
| S962K | F: CAAGCACCTATGAAAACCCTCAACACG<br>R: CGTGTTGAGGGTTTTCATAGGTGCTTG |
| T963K | F: GCACCTATGTCAAAGCTCAACACGTTGAAAG<br>R: CTTTCAACGTGTTGAGCTTTGACATAGGTGC |
| L964K | F: GCACCTATGTCAACCAAGAACACGTTGAAAGC<br>R: GCTTTCAACGTGTTCTTGGTTGACATAGGTGC |
| N965K | F: GTCAACCCTCAAGACGTTGAAAGCAG<br>R: CTGCTTTCAACGTCTTGAGGGTTGAC |
| T966K | F: CAACCCTCAACAAGTTGAAAGCAGG<br>R: CCTGCTTTCAACTTGTTGAGGGTTG |
| L967K | F: CCCTCAACACGAAGAAAGCAGGTGGG<br>R: CCCACCTGCTTTCTTCGTGTTGAGGG |
| A969K | F: CCTCAACACGTTGAAAAAAGGTGGGATTC<br>R: GAATCCCACCTTTTTTCAACGTGTTGAGG |
| G970K | F: CAACACGTTGAAAGCAAAGGGGATTCTTAATAGATTC<br>R:GAATCTATTAAGAATCCCCTTTGCTTTCAACGTGTTG |
| S1058K | F: CACTCATCTTGTTACAAAGTTAAAAGGACTATGG<br>R: CCATAGTCCTTTTAACTTTGTAACAAGATGAGTG |
| L1059K | F: ACAAGCAAAAAAGGACTATGGACACTTCGTG<br>R: GTCCTTTTTTGCTTGTAACAAGATGAGTGAAAATTGG |
| K1060R | F: AGAGGACTATGGACACTTCGTGCCTTCG<br>R: GTGTCCATAGTCCTCTTAAGCTTGTAACAAGATG |
| G1061K | F: GTTACAAGCTTAAAAAAGCTATGGACACTTCG<br>R: CGAAGTGTCCATAGCTTTTTTAAGCTTGTAAC |
| L1062K | F: CAAGCTTAAAAGGAAAATGGACACTTCGTG<br>R: CACGAAGTGTCCATTTTCCTTTTAAGCTTG |
| W1063K | F: GCTTAAAAGGACTAAAGACACTTCGTG<br>R: CACGAAGTGTCTTTAGTCCTTTTAAGC |
| T1064K | F: AAGGACTATGGAAACTTCGTGCCTTC<br>R: GAAGGCACGAAGTTTCCATAGTCCTT |
| L1065K | F: GGACTATGGACAAAGCGTGCCTTCGGACG<br>R: CGTCCGAAGGCACGCTTTGTCCATAGTCC |
| N901K | F: AAAAGCTATGCAGTGATTATCACCAGCACC<br>R: CACTGCATAGCTTTTATTTCTACTATGAGTACTATTCC |
| S902K | F: ACAAATATGCAGTGATTATCACCAGCACCAG<br>R: TCACTGCATATTTGTTATTTCTACTATGAGTACTATTCC |
| Y903K | F: CAGCAAAGCAGTGATTATCACCAGCACC<br>R: GATAATCACTGCTTTGCTGTTATTTCTACTATGAGTAC |
| A904K | F: ACAGCTATAAAGTGATTATCACCAGCACCAG<br>R: TCACTTTATAGCTGTTATTTCTACTATGAGTACTATTCC |
| V905K | F: CAGCTATGCAAAGATTATCACCAGCACCAG<br>R: ATCTTTGCATAGCTGTTATTTCTACTATGAGTACTATTC |
| T908K | F: GCAGTGATTATCAAAAGCACCAGTTCGTATTATG<br>R: GTGCTTTTGATAATCACTGCATAGCTGTTATTTCTAC |
| S909K | F: GCAGTGATTATCACCAAAACCAGTTCGTATTATG<br>R: GGTGATAATCACTGCATAGCTGTTATTTCTACTATG |
| T910K | F: CAGTGATTATCACCAGCAAAAGTTCGTATTATGTG<br>R: GCTGGTGATAATCACTGCATAGCTGTTATTTC |

| | |
|---|---|
| S911K | F: CCAGCACCAAATCGTATTATGTGTTTTACATTTACG<br>R: AATACGATTTGGTGCTGGTGATAATCACTGC |
| S912K | F: CCAGCACCAGTAAGTATTATGTGTTTTACATTTACG<br>R: AATACTTACTGGTGCTGGTGATAATCACTGC |
| D565A | F: GAGCAGTATACAAAGCTGCTGATTTGTA<br>R: CAATCAGCAGCTTTGTATACTGCTC |
| D567A | F: GCAGTATACAAAGATGCTGCTTTGTATTTATTAGAC<br>R: GTCTAATAAATACAAAGCAGCATCTTTGTATACTGC |
| D565A/D567A | F: GCAGTATACAAAGCTGCTGCTTTGTATTTATTAGACTCTCC<br>R: GCAGTATACAAAGCTGCTGCTTTGTATTTATTAGACTCTCC |
| D565G | F: GCAAGAGCAGTATACAAAGGTGCTGATTTGTATTTATTAG<br>R: CTAATAAATACAAATCAGCACCTTTGTATACTGCTCTTGC |

**Table S7. Conditions for immunoprecipitation**

| Antibody | Mab/Pab | Species | Protein A/G | Incubation 4°C | Wash Buffer |
|---|---|---|---|---|---|
| MM13-4 | Mab | Mouse | G | overnight | PBS, 0.2% SDS, 1% Triton X-100 |
| E1-22 | Pab | Rabbit | A | 3 h | 10 mM Tris-HCl pH 8.6, 300 mM NaCl, 0.05% SDS, 0.05% Triton X-100 |
| TMD1C | Pab | Rabbit | A | 3 h | 10 mM Tris-HCl pH 8.6, 300 mM NaCl, 0.05% SDS, 0.05% Triton X-100 |
| 3G11 | Mab | Rat | G | overnight | 10 mM Tris-HCl pH 8.6, 300 mM NaCl, 0.05% SDS, 0.05% Triton X-100 |
| MrPink | Pab | Rabbit | A | overnight | 10 mM Tris-HCl pH 8.6, 300 mM NaCl, 0.1% SDS, 0.05% Triton X-100 |
| G449/G450 | Pab | Rabbit | A | overnight | 10 mM Tris-HCl pH 8.6, 300 mM NaCl, 0.1% SDS, 0.05% Triton X-100 |
| 570 | Mab | Mouse | A | overnight | 50 mM Tris-HCl pH 8.0, 150 mM NaCl, 1 mM EDTA |
| 217 | Mab | Mouse | A | overnight | 50 mM Tris-HCl pH 8.0, 150 mM NaCl, 1 mM EDTA |
| I4N | Pab | Rabbit | A | overnight | 50 mM Tris-HCl pH 8.0, 150 mM NaCl, 1 mM EDTA |
| TMD2C | Pab | Rabbit | A | 3 h | 50 mM Tris-HCl pH 8.0, 150 mM NaCl, 1 mM EDTA |
| 596 | Mab | Mouse | A | 3 h | 20 mM MES, 100 mM NaCl, 30 mM Tris-HCl pH 7.4) with 0.5% Triton X-100 |
| 2-39.14 | Mab | Mouse | G | 3 h | 20 mM MES, 100 mM NaCl, 30 mM Tris-HCl pH 7.4) with 0.5% Triton X-100 |
| 2-3.5 | Mab | Mouse | G | 3 h | 20 mM MES, 100 mM NaCl, 30 mM Tris-HCl pH 7.4) with 0.5% Triton X-100 |
