## Supplementary figures and images for "ABC-transporter CFTR folds with high fidelity through a modular, stepwise pathway"

### Suppl. Figures

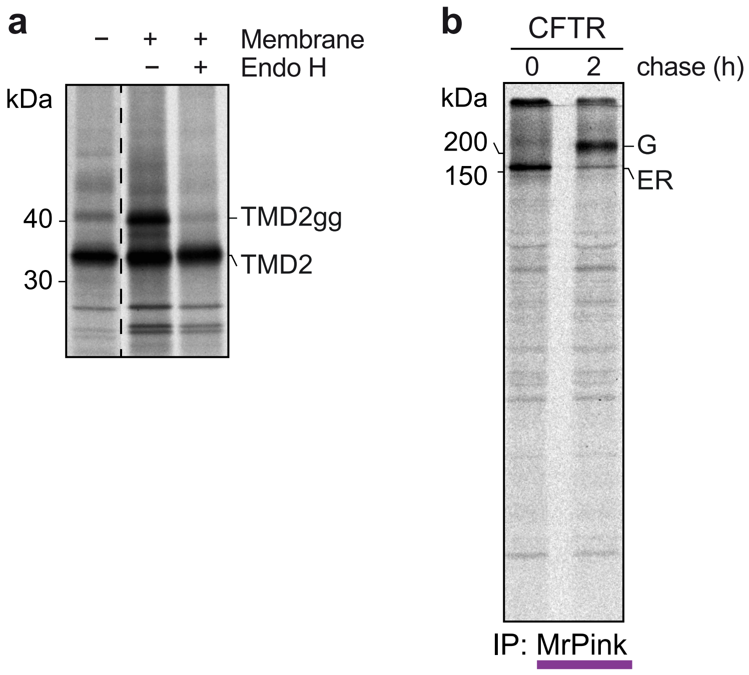

Figure S2

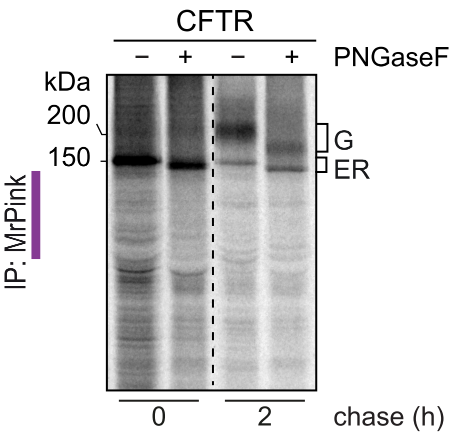

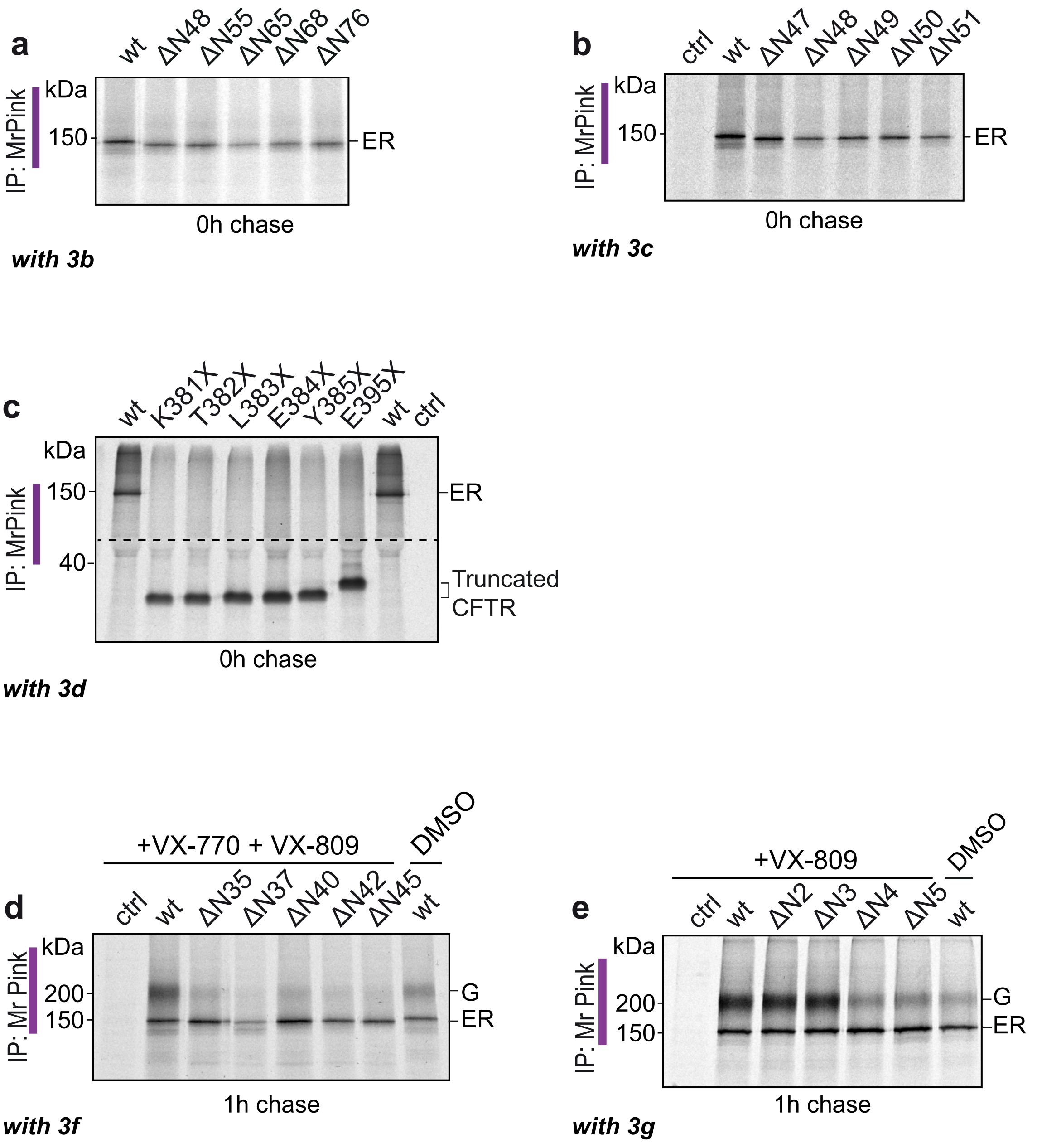

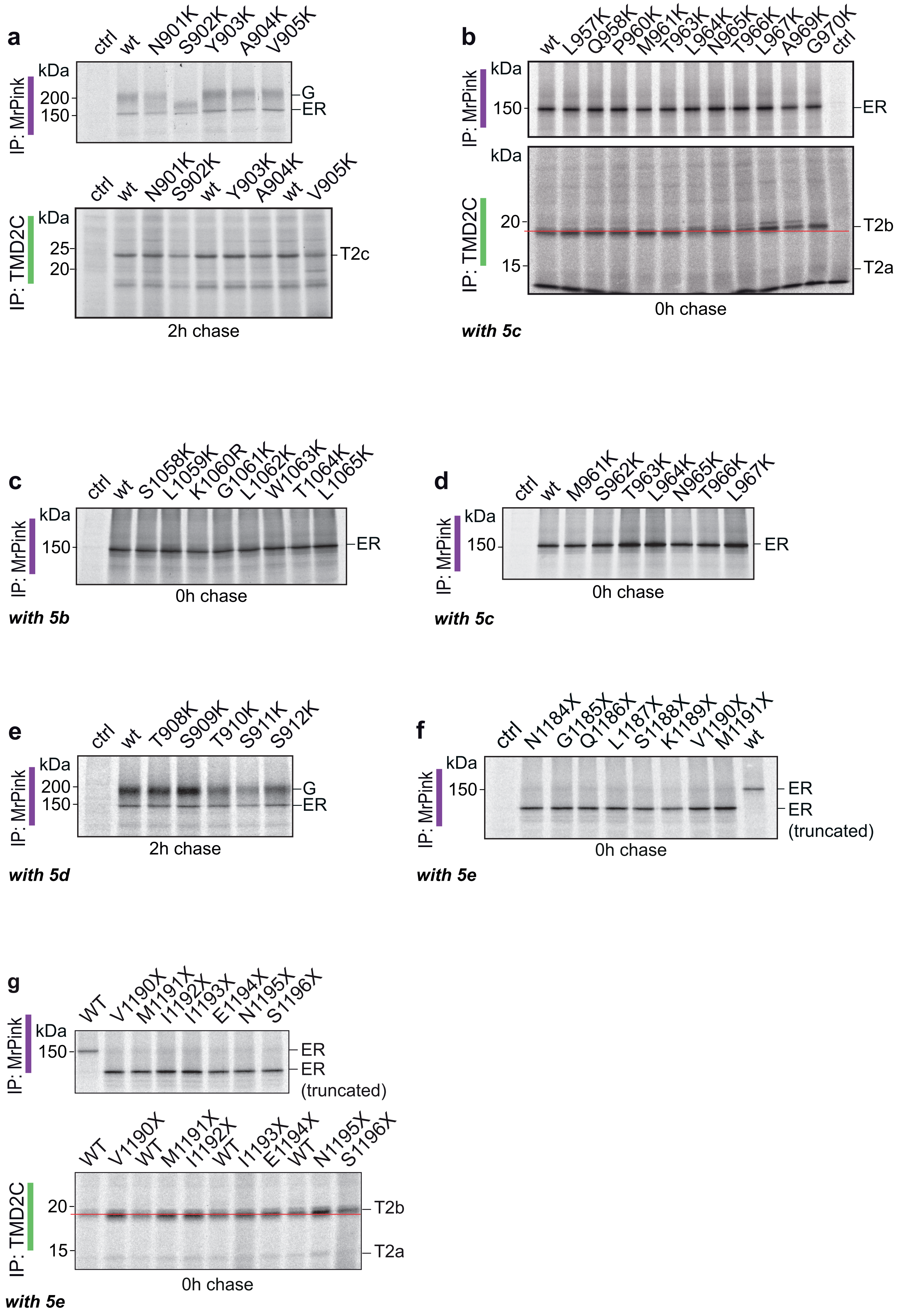
